# A Comparison of mRNA Sequencing with Random Primed and 3’-Directed Libraries

**DOI:** 10.1101/098905

**Authors:** Yuguang Xiong, Magali Soumillon, Jie Wu, Jens Hansen, Bin Hu, Johan G.C. van Hasselt, Gomathi Jayaraman, Ryan Lim, Mehdi Bouhaddou, Loren Ornelas, Jim Bochicchio, Lindsay Lenaeus, Jennifer Stocksdale, Jaehee Shim, Emilda Gomez, Dhruv Sareen, Clive Svendsen, Leslie M. Thompson, Milind Mahajan, Ravi Iyengar, Eric A. Sobie, Evren U. Azeloglu, Marc R. Birtwistle

**Affiliations:** Department of Pharmacological Sciences and DToxS LINCS Center, Icahn School of Medicine at Mount Sinai, New York, NY; Broad Institute of MIT and Harvard, Cambridge, MA; Department of Biological Chemistry, University of California, Irvine, CA USA; UCI MIND, University of California, Irvine, CA USA; Board of Governors-Regenerative Medicine Institute, Cedars-Sinai Medical Center, Los Angeles, CA, USA; iPSC Core, The David and Janet Polak Foundation Stem Cell Core Laboratory, Los Angeles, CA, USA; Department of Biomedical Sciences, Cedars-Sinai Medical Center, Los Angeles, CA, USA; Department of Psychiatry and Human Behavior, Neurobiology and Behavior, University of California, Irvine, CA USA; Department of Genetics, Icahn School of Medicine at Mount Sinai, New York, NY 10029

## Abstract

Deep mRNA sequencing (mRNAseq) is the state-of-the-art for whole transcriptome measurements. A key step is creating a library of cDNA sequencing fragments from RNA. This is generally done by random priming, creating multiple sequencing fragments along the length of each transcript. A 3’ end-focused library approach cannot detect differential splicing, but has potentially higher throughput at lower cost (~10-fold lower), along with the ability to improve quantification by using transcript molecule counting with unique molecular identifiers (UMI) to correct for PCR bias. Here, we compare implementation of such a 3’-digital gene expression (3’-DGE) approach with “conventional” random primed mRNAseq, which has not yet been done. We find that while conventional mRNAseq detects ~15% more genes, the resulting lists of differentially expressed genes and therefore biological conclusions and gene signatures are highly concordant between the two techniques. We also find good quantitative agreement on the level of individual genes between the two techniques in terms of both read counts and fold change between two conditions. We conclude that for high-throughput applications, the potential cost savings associated with the 3’-DGE approach are a very reasonable tradeoff for modest reduction in sensitivity and inability to observe alternative splicing, and should enable much larger scale studies focused on not only differential expression analysis, but also quantitative transcriptome profiling. The computational scripts and programs, along with experimental standard operating procedures used in our pipeline presented here, are freely available on our website (www.dtoxs.org).

## Introduction

Massively parallel sequencing of mRNA, or mRNAseq, was first introduced in 2008 (Mortazavi et al. 2008; Trapnell et al. 2010, 2009), and since has rapidly become the preferred method for whole transcriptome measurements (Li et al. 2014a; Conesa et al. 2016; Li et al. 2014b; Risso et al. 2014; Wang et al. 2014; Su et al. 2014; Byron et al. 2016; Trapnell et al. 2013), culminating recently with the announcement by Illumina of the discontinuation of the Human Expression Array BeadChip (HumanHT-12 v4 as of 9 Dec 2016). The procedure consists of two basic steps. First is library preparation, which consists of converting mRNA isolated from samples into cDNA that is compatible with the deep sequencing platform. Next is the sequencing itself, which often consists of paired-end protocols on the Illumina HiSeq platform to generate millions of reads per sample. The resulting quantification of transcript levels comes from counting the reads aligned to each transcript, followed with normalizing by transcript length (since # of reads depends on transcript length due to random priming), and the total number of reads (sequencing depth) (Trapnell et al. 2013; Conesa et al. 2016). Analysis of alternative splicing is also possible with paired-end sequencing, which, along with a reference transcriptome, puts strong constraints on the distance between reads and allows assessment of compatibility with known splice isoforms (Conesa et al. 2016; Trapnell et al. 2010).

Although the data resulting from mRNAseq is often considered superior to that obtained by the former transcriptome measurement standard, the microarray, the cost per sample is currently ~$300–$500 (depending on various options), precluding widespread high-throughput application. The primary component of this cost, typically ~70–90% depending on core facility and desired sequencing depth, is the library preparation. Thus, reduction in library preparation cost is expected to pay large returns for ability to increase mRNAseq throughput.

Another current issue with mRNAseq data are the biases introduced by PCR during library preparation. Because every sequence has a potentially different propensity to be amplified during PCR, the resulting quantitative representation of transcripts in the sequencing library is non-linearly distorted from the original abundance in ways that are difficult to predict. One way that has been shown to correct for this bias is to tag every transcript molecule with a unique random nucleotide sequence prior to amplification (Kivioja et al. 2011; Islam et al. 2014; Smith and Birtwistle 2016). Such sequences are termed unique molecular identifiers (UMIs). UMIs allow removal of PCR bias by only counting reads for a gene that have different UMI sequences, and ignoring those that have the same UMI sequence, since they came from the same original transcript molecule and thus are PCR duplicates. For this reason, such quantification approaches are also often referred to as transcript counting or digital gene expression (DGE).

Here, we demonstrate the use of a library preparation method that takes advantage of 3’-end creation of cDNA (poly-T priming) and incorporates UMI-based quantification, but only costs ~$30 per sample, greatly reducing total mRNAseq cost to ~$50 per sample, a nearly 10fold reduction in typical cost. Because it is not yet clear to what extent the results from this 3’-DGE library preparation method overlap with the current gold-standard of random primed conventional mRNA sequencing, we performed a comprehensive comparison of data obtained by both methods, from the same RNA samples. We find that 3’-DGE has only about ~15% lower sensitivity than conventional random primed mRNAseq, good quantitative agreement, and high overlap in lists of differentially expressed genes. We conclude that the 3’-DGE approach used here is a viable low-cost alternative to conventional random-primed mRNAseq for high-throughput applications, particularly when looking for differentially expressed genes between treatment conditions, as is a common goal for transcriptomic studies, but also for simple expression profiling which is becoming more commonplace in single-cell mRNAseq (DeLaughter et al. 2016; Wagner et al. 2016; La Manno et al. 2016; Olsson et al. 2016) or tissue-level examinations (Ardlie et al. 2015; Melé et al. 2015; Genomics et al. 2015).

## Results and Discussion

### 3’-Digital Gene Expression Versus Conventional mRNA Sequencing

Both conventional and 3’-DGE mRNAseq consist of (i) library preparation, (ii) sequencing, (iii) alignment to a reference genome and (iv) quantification, but there are differences between the two approaches in each of these four main steps (Fig. 1 illustrates some of these). For conventional library preparation, following isolation of mRNA from total RNA (in our case with oligo dT beads—see Methods), cDNA is typically synthesized via random priming of thermally-sheared mRNA. In 3’-DGE, cDNA is generated via 3’-directed oligo dT priming, during which unique molecular identifiers (UMI) are incorporated (for quantification purposes—see below), and strand-specificity is preserved. For conventional sequencing, single-end or paired-end constant read length is used (we used 100 bp single-end in this paper). In 3’-DGE, paired-end is required. At least 16 bp are needed on the first read to capture UMI and sample barcode data. We used 46 bp on the second read to acquire transcript-specific sequence data. Focus on the 3’ end of the transcript coupled with lack of transcript-specific information on both reads precludes identification of alternative splicing for 3’-DGE. For conventional alignment, the entire reference transcriptome is used since sequenced fragments are randomly distributed. In 3’-DGE, a reduced transcriptome is used, since sequenced fragments are localized near the 3’ end of transcripts. Because of this, there are slight differences in the gene lists obtained from each alignment (Table S1), but a large majority of protein-coding genes (22,811) are shared between the approaches. For conventional quantification, the total number of uniquely aligned reads to a particular gene is used, termed read counts. Read counts are often divided by the average transcript length and read depth (units of **r**eads **p**er **k**ilobase of transcript length per **m**illion mapped reads—RPKM). In 3’-DGE, the total number of uniquely aligned reads to particular genes is available, but is only an intermediate to the final quantification by UMI counts for each gene. This UMI count metric corrects for PCR bias by removing reads that align to the same genomic region and share the same UMI sequence (this strongly suggests they arose from the same original transcript molecule).

**Figure 1.**
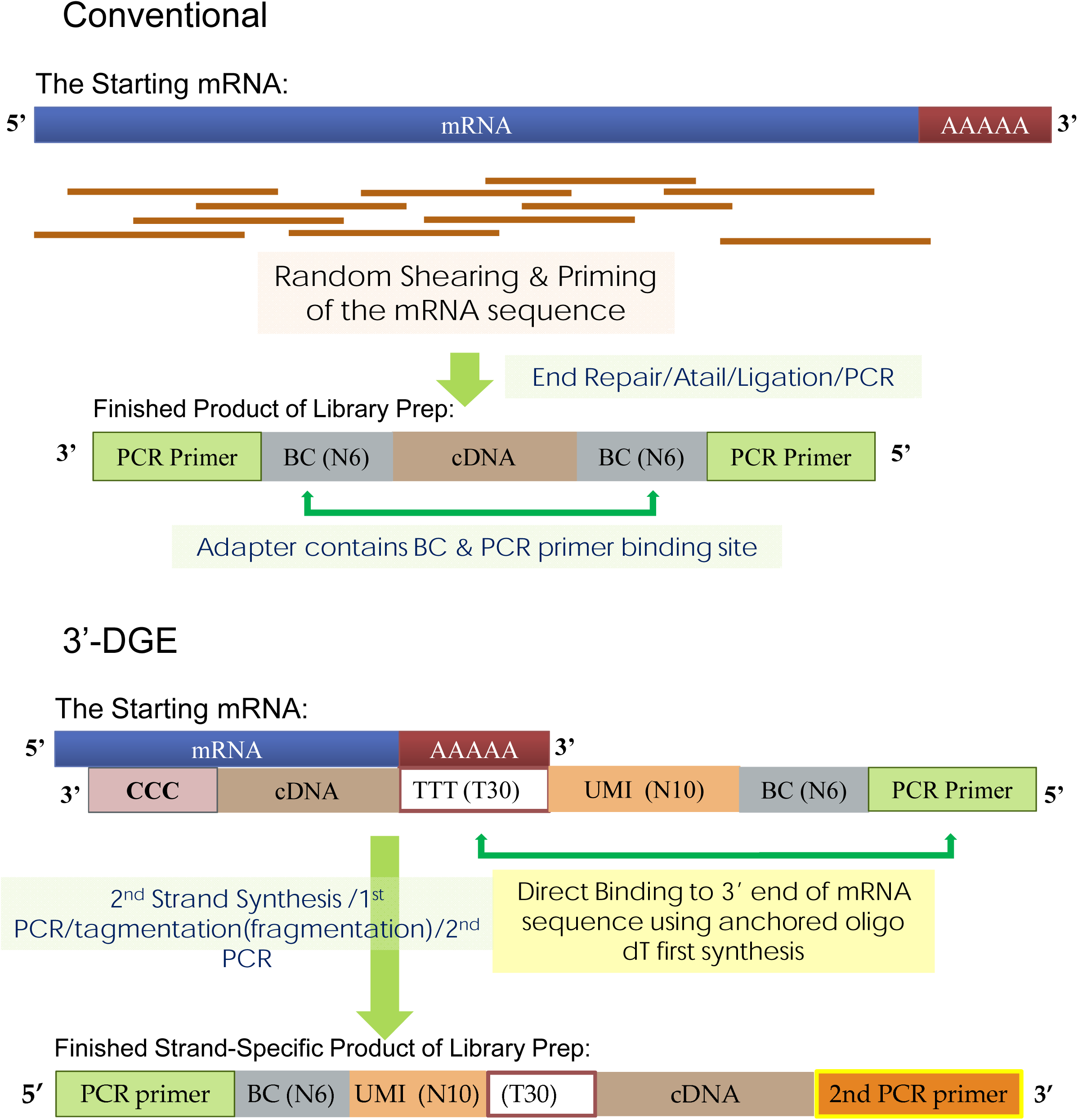
Schematic of Library Construction Differences between Conventional and 3’-DGE mRNA Sequencing. BC: Barcode; UMI: Unique Molecular Identifier.

To gain a thorough practical understanding of the differences between data obtained by 3’-DGE and conventional mRNAseq, we isolated RNA from 16 PromoCell primary cardiomyocyte cultures treated with either DMSO vehicle control (eight biological replicates), sorafenib (four biological replicates) or sunitinib (four biological replicates) for 48 hours. These experiments were part of a larger signature generation project in our DToxS LINCS center focused on cellular signatures for cardiotoxicity of kinase inhibitors (www.dtoxs.org and www.lincsproject.org) We expected, based on our prior data, that sorafenib would induce large changes in gene expression, whereas sunitinib would induce negligible changes in gene expression, providing positive and negative control test cases for differential expression analysis. The 16 RNA samples were analyzed for quality, and then split and sent to either the conventional or 3’-DGE mRNAseq pipeline for library preparation, sequencing, alignment and quantification (see Methods and Fig. S1). The average read depth for conventional was 5.9×10^6^ reads / sample, and for 3’-DGE was 3.8×10^6^ reads / sample. Read depth was also consistent across samples (Fig. S2), and overall read count distribution was similar for the two techniques (Fig. S3).

### Fidelity of Sequence Alignments for 3’-DGE

Because sequence information in 3’-DGE comes from a restricted region of the exome that may have reduced sequence entropy amongst genes, we first investigated the fidelity of sequence alignments. For each gene with at least four read counts (summed across all samples), we quantified the proportion of reads that align only to that gene, and looked at the frequency distribution of this proportion across genes (Fig. 2). The distribution is highly skewed towards proportions > 0.95, indicating most genes are quantified by reads that align only to that single gene. This proportion is less than 0.10 for a small number of genes (738 out of 14,574; gene names are in Table S2). Thus these 738 genes are not able to be reliably quantified without further assumptions and considerations. The reads associated with these unquantifiable genes account for most such degenerately-aligned reads (>50%). We conclude from these data that despite the fact that reads are 3’ end-focused, a large and sufficient majority can be reliably mapped to individual genes with high fidelity. This feature may be facilitated by the strand-specificity of the 3’-DGE library preparation method. From this point forward we only consider read counts that are reliably aligned to a single gene.

**Figure 2.**
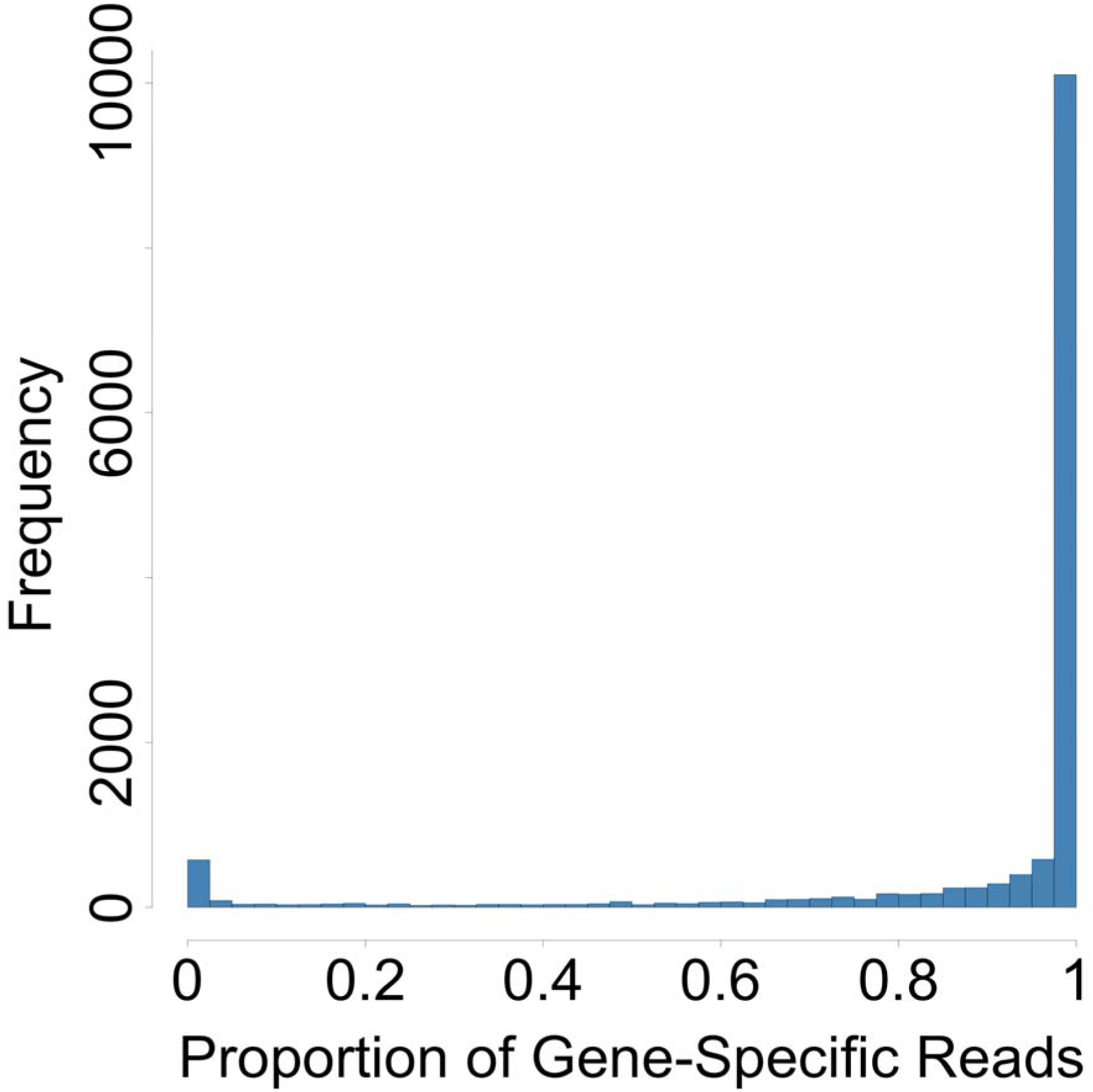
Fidelity of Sequence Alignments in 3’-end Digital Gene Expression (3’-DGE) Method. For each of the 14,574 genes with greater than four counts (summed across all 16 samples), the proportion of reads aligned to only that one gene was calculated. This proportion is close to 1 for most genes, indicating reliable quantification.

### Sensitivity to Detect Gene Expression as a Function of Read Depth

We wanted to determine the sensitivity of the two techniques to detect expression of the 22,811 genes shared between the reference sequence databases. To do this we employed a read removal approach, where each sample’s dataset was progressively downsized (Fig. 3A and Figs. S4–5). Read counts were removed from each gene with probability proportional to that gene’s overall representation in the dataset, and expression of a gene was considered detected with four or greater reads (see Methods). We verified that this stochastic nature of read removal introduced negligible variability into our results (Fig. S6).

**Figure 3A.**
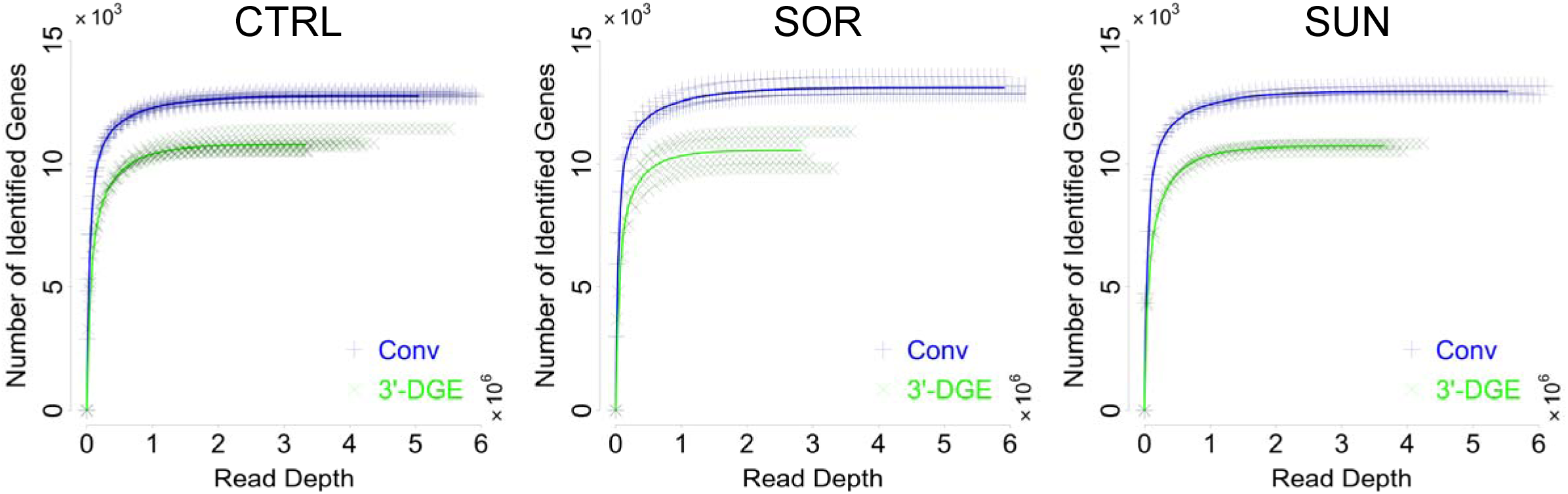
Sensitivity of Conventional (Conv) and 3’-end Digital Gene Expression (3’-DGE) mRNA Sequencing Methods. Gene-wise reads are removed from every sample in a probability proportional to the abundance of the gene in a sample, to generate a set of the number of identified genes over a range of simulated read depths. The curves for individual replicate samples are shown with the thinner points, showing in general low variability. The average is shown with the solid line.

In all treatment cases, we observed a hyperbolic dependency between read depth and number of detected genes. We parameterized these relationships with a Michaelis-Menten model to quantify differences between the techniques (Table 1). We found that the estimated parameter values were consistent across samples for each measurement type. The *V_max_* parameter indicates that on average conventional mRNAseq detects approximately 15% (~2000) more genes. The genes that were differentially detected by conventional were very consistent across treatments, and were greater in number than those differentially detected by 3’-DGE as expected (Fig. 3B). The overall fraction of genes identified by only conventional or 3’-DGE was small, indicating good concordance between the techniques (>10^4^ vs. <10^3^). The *K_m_* parameter indicates that on average conventional mRNAseq is slightly more sensitive; however, at read depths even far below typical levels (10^6^ / sample), both techniques detect ~95% of the maximum possible. We conclude that conventional mRNAseq detects the expression of approximately 15% more genes (~13,000 vs. ~11,000), but both techniques saturate in terms of detecting new genes at greater than 2-3 million reads / sample. It should be noted here that the 3’-DGE method was designed to be compatible with very little total RNA extract (e.g. single cell RNAseq), so this may well account for the differences observed here. Also, detection is strongly distinct from statistical power, so 2-3 million reads / sample my not be sufficient to detect differential expression of lowly expressed genes between two conditions, despite being detected.

**Figure 3B.**
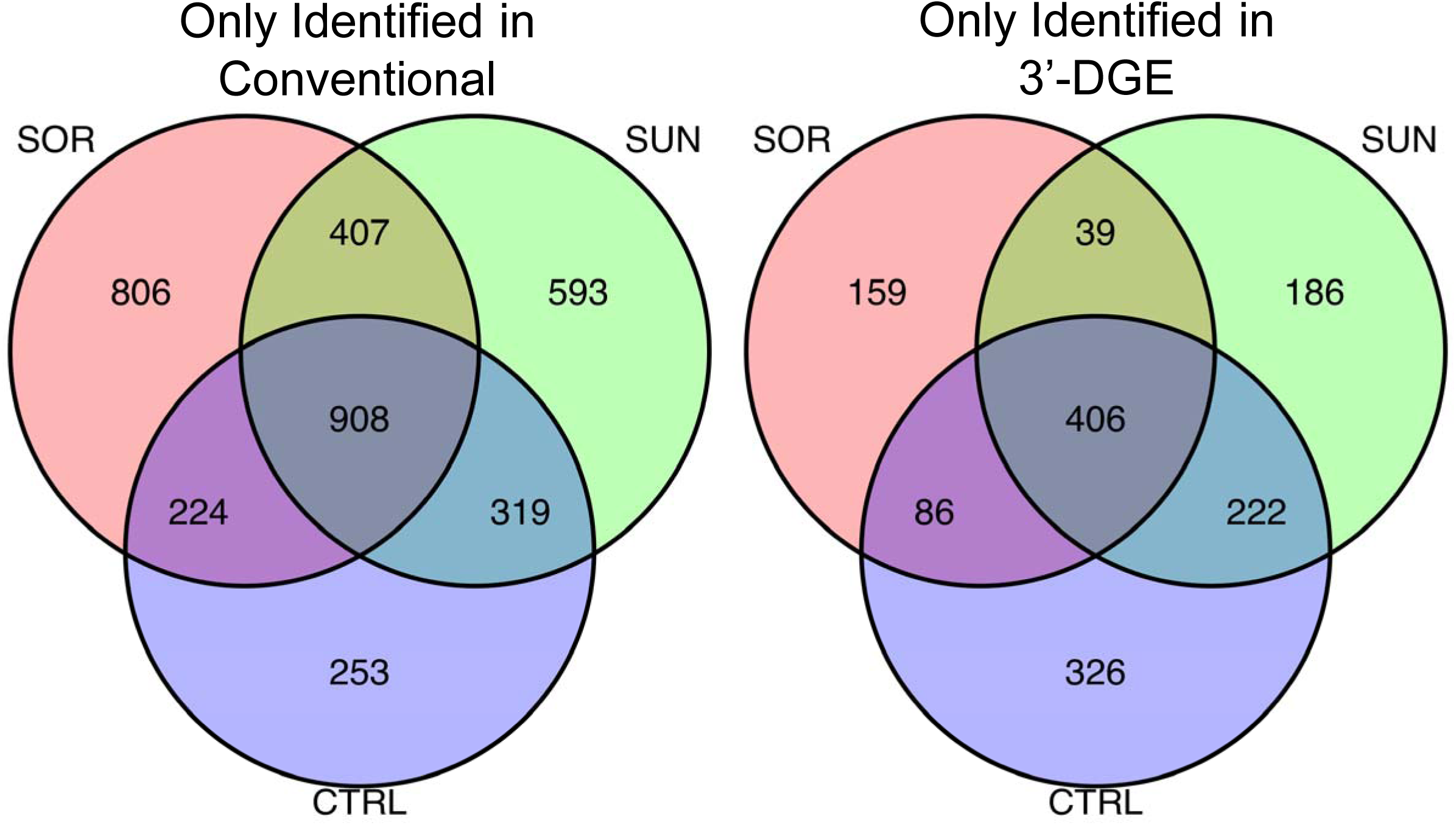
Genes Identified in Conventional but not 3’-DGE, and Vice Versa. Each replicate from both mRNAseq technologies were downsampled via read removal to a common read depth (2.8 million reads per sample), and the differences in identified genes were analyzed. Most genes identified in conventional but not 3’-DGE were shared across treatment conditions, and likewise for those identified by 3’-DGE but not conventional.

**Table 1.** The Best-Fit Parameter Values of a Michaelis-Menten Model to the Average Curves in Fig.3A. The function *drm* in the R package *drc*, using the fit function *mm*, was used.

| Technology | Treatment | $K_m$ (reads) | $V_{max}$ (genes) |
| --- | --- | --- | --- |
| Conv | CTRL | 49,719 | 12,905 |
| Conv | SOR | 45,441 | 13,202 |
| Conv | SUN | 50,902 | 13,099 |
| 3'-DGE | CTRL | 75,277 | 11,117 |
| 3'-DGE | SOR | 60,979 | 10,884 |
| 3'-DGE | SUN | 70,673 | 11,006 |

### Quantitative Comparison Between the Techniques

We next wanted to do a quantitative comparison of expression gene-by-gene for matched samples and treatment conditions. The first step in doing so is to normalize read depth amongst all the samples. To do that we used the read removal process as above to downsample each dataset to the lowest common read depth, which was ~2.8 million reads per sample. Correlation amongst replicates within the same technique was very high (Fig. 4A), but the resulting x-y scatterplots of conventional read counts vs. DGE read counts yielded poor correlation (Fig. 4B). We reasoned that this poor correlation could be due to the fact that conventional read counts depend strongly on transcript length, whereas in principal those from 3’-DGE do not. When we normalized conventional read counts by transcript length, the correlation improved significantly (Fig. 4C). We observed similar agreement on the level of individual sample-to-sample correlations (Fig. S7). We conclude that the two techniques show reasonable quantitative agreement with one another.

**Figure 4A.**
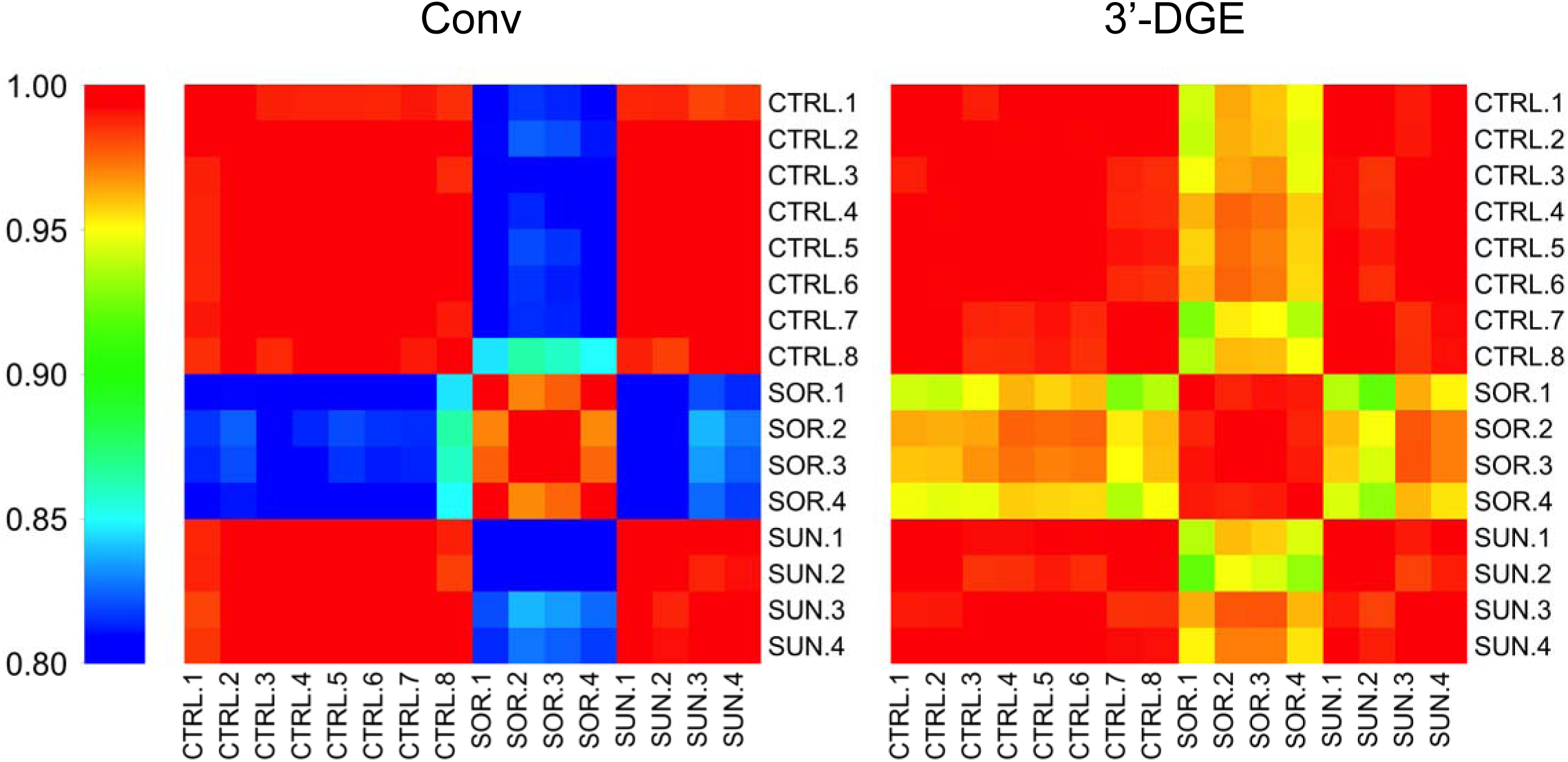
Correlations of the Replicate Samples from the Conventional (Conv) Read Counts and 3’-end Digital Gene Expression (3’-DGE) Read Counts. The replicate samples obtained by the same method correlate well with each other at each condition. Control (CTRL), Sorafenib (SOR), Sunitinib (SUN).

**Figure 4B-C.**
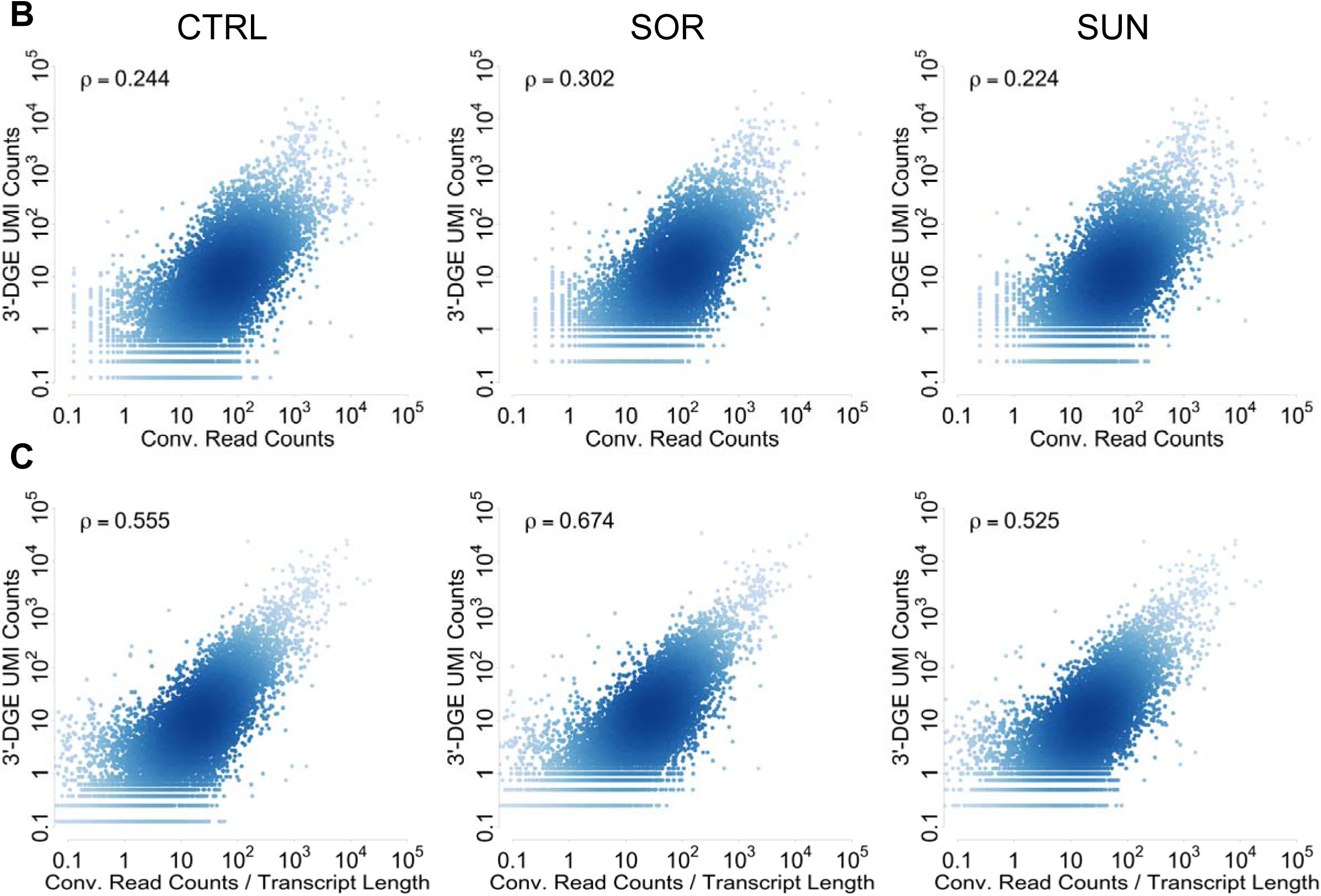
Quantitative Gene-wise Comparison between 3’-end Digital Gene Expression (3’-DGE) and Conventional Techniques. Datasets are downsampled to a common read depth of 2.8 million reads, and then gene-by-gene comparisons are made via scatter plots. To generate a reduced UMI count dataset, upon removal of a read count, UMI counts were removed with probability proportional to the ratio between UMI counts and read counts for that gene (accounting for PCR bias). Density of points in scatter plots is indicated by depth of color. Inset text box shows Pearson correlation. In all plots, data are scaled so units are comparable. **B.** Scatterplots of UMI counts for 3’-DGE versus read counts for conventional, without normalization by average transcript length. There is a general trend of agreement but correlation is low for quantitative agreement. **C.** Scatterplots of UMI counts for 3’-DGE versus transcript length-normalized read counts for conventional. Quantitative agreement is significantly improved upon this normalization.

We also used such scatter plot analysis to yield insight into potential biases in detection for each technique (Fig. 4D). We first took the average read count values for each gene across all 16 samples, with the thinking that strong biases should largely be sample independent. As a simple and conservative approach, we drew lines parallel to x=y that flank the typical variance in the data, and identified points falling outside of this range (Fig. 4D). We identified a small relative number of such genes—165 for 3’-DGE and 98 for conventional (Tables S5-S6). These genes should be interpreted with caution when appearing as differentially expressed in downstream analyses.

**Figure 4D.**
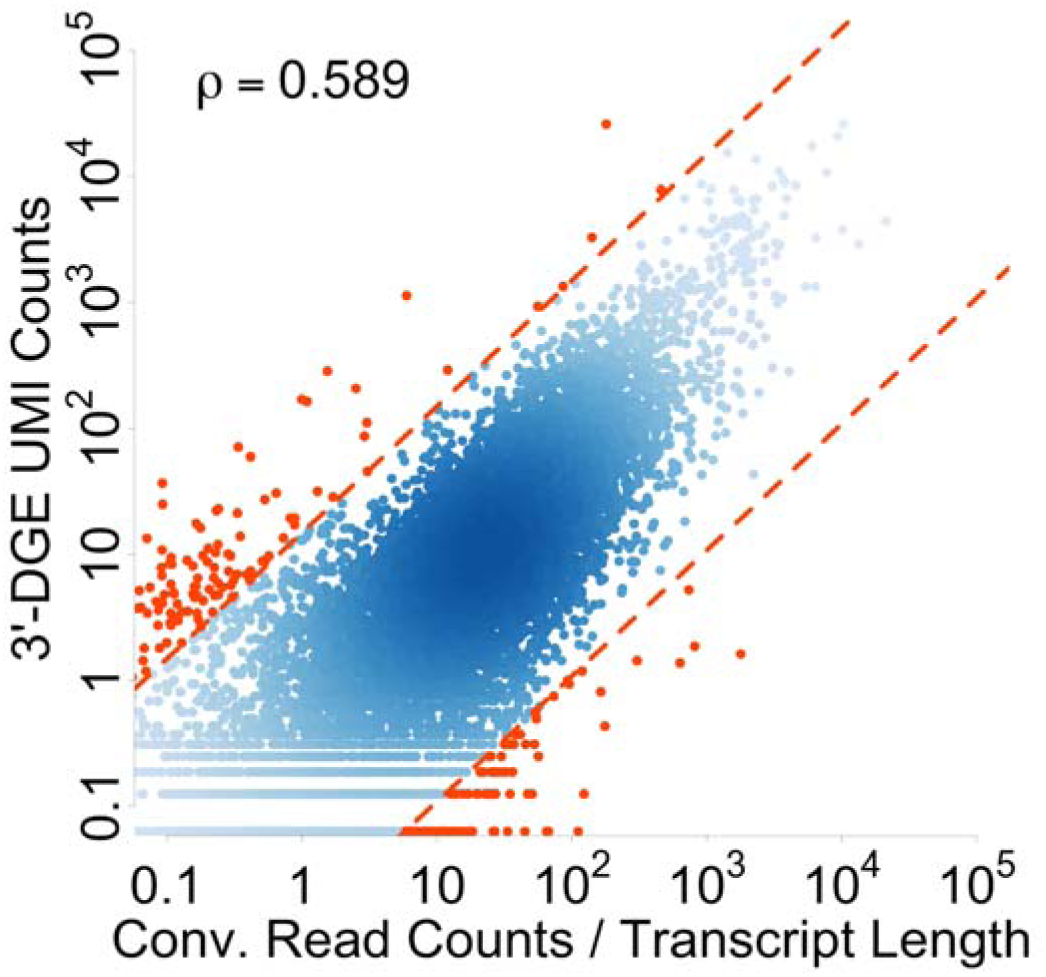
Potential Biases of 3’-end Digital Gene Expression (3’-DGE) or Conventional Techniques. We averaged data from all 16 read depth-normalized samples and defined lines that flank the typical variance in the data to identify genes that have evidence of bias in quantification. Genes falling outside of this range are reported in Tables S5–6.

### Differential Expression Analysis

The typical endpoint of an mRNA sequencing experiment is testing for statistical significance of differential expression between two or more conditions. There are many software packages available for doing this; here we used EdgeR (Robinson et al. 2009) (see Methods for specific software settings). Specifically, we compared DMSO vehicle-treated control (CTRL) to either sunitinib (SUN) or sorafenib (SOR) treatment. A typical definition of a “differentially expressed gene” is that which has a false discovery rate (FDR) of lower than 0.1 (Fig. 5A). Both conventional and 3’-DGE data show that sunitinib causes very few genes to be differentially expressed, whereas sorafenib causes differential expression of several thousand genes (Fig. 5A). For the 3,136 sorafenib-associated gene expression changes identified by both conventional and 3’-DGE, there is a strong correlation between the p-values for differential expression (Fig. 5B). However, there are another ~3,600 genes identified as differentially expressed by conventional that are not found by 3’-DGE. The reason for this is a simple statistical power argument due to UMI counting versus read counting. While using UMI counts as the quantification metric in principal improves precision due to removal of PCR bias, at the same time, it reduces effective read depth and therefore statistical power. We thus decided to forgo strict FDR cutoffs and instead analyze trends visible from all 10,121 genes with defined p-values for differential expression. An expected cone-shape that falls along the x=y line is observed as statistical significance varies from weak to strong, consistent with the above hypothesis and further suggestive of concordance between the techniques (Fig. 5C). This concordance is also evident from the strong agreement between the log_2_-fold changes for differential expression from each technique (Fig. 5D). This strongly suggests the biological conclusions drawn from either technique will be concordant. Lastly, we ranked the entire list of 22,811 genes for each mRNAseq technique and drug, and then analyzed their overlap via the rank-rank hypergeometric test (Fig. 5E-F). Each spot in the heatmap quantifies the statistical significance of overlap between the genes in the ranked list up to that point, starting from the top left corner, via Fisher’s exact test. To distinguish between up and down regulation, we signed the p-value accordingly, placing down-regulated genes with high statistical significance at the bottom of the list (Fig. 5E). As expected, the ranked lists for sorafenib-treated samples from conventional and 3’-DGE show large agreement, as indicated by the red color down the diagonal. The ranked lists for suninitib-treated samples from conventional and 3’-DGE show much less agreement, also as expected for gene lists with little to no differential expression. Likewise, cross comparison between treatments and sequencing type also yield little agreement as expected. We conclude that conventional and 3’-DGE mRNAseq techniques generate highly similar signatures of gene expression for both up and down-regulated genes.

**Figure 5A.**
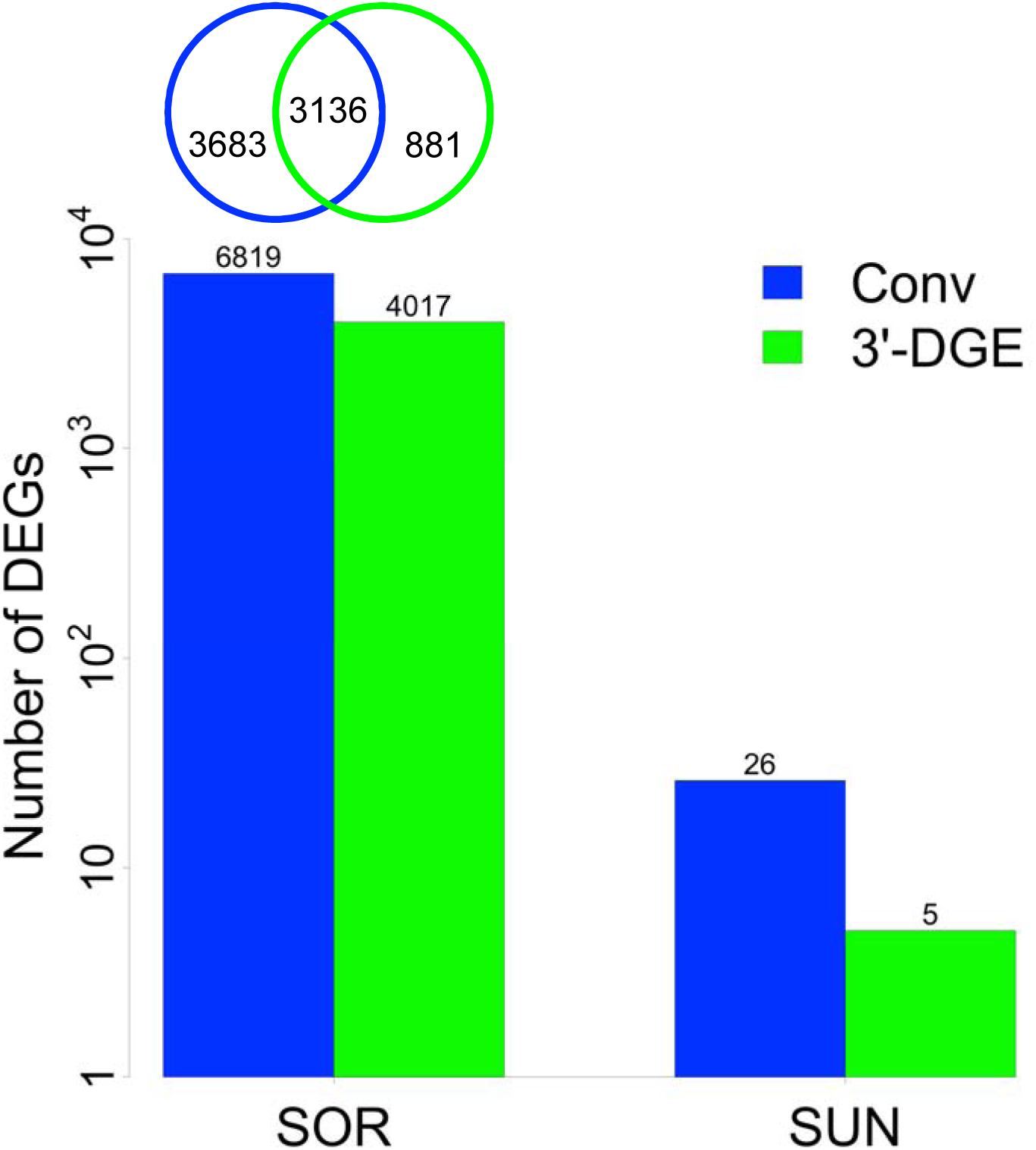
Differential Expression Analysis. Control (CTRL) data are compared Sorafenib (SOR) or Sunitinib (SUN) to identify differentially expressed genes (DEGs) using EdgeR for both conventional (Conv) and 3’-end digital gene expression (3’-DGE) datasets. A gene is defined as differentially expressed using a false discovery rate (FDR) cutoff of 0.1.

**Figure 5B.**
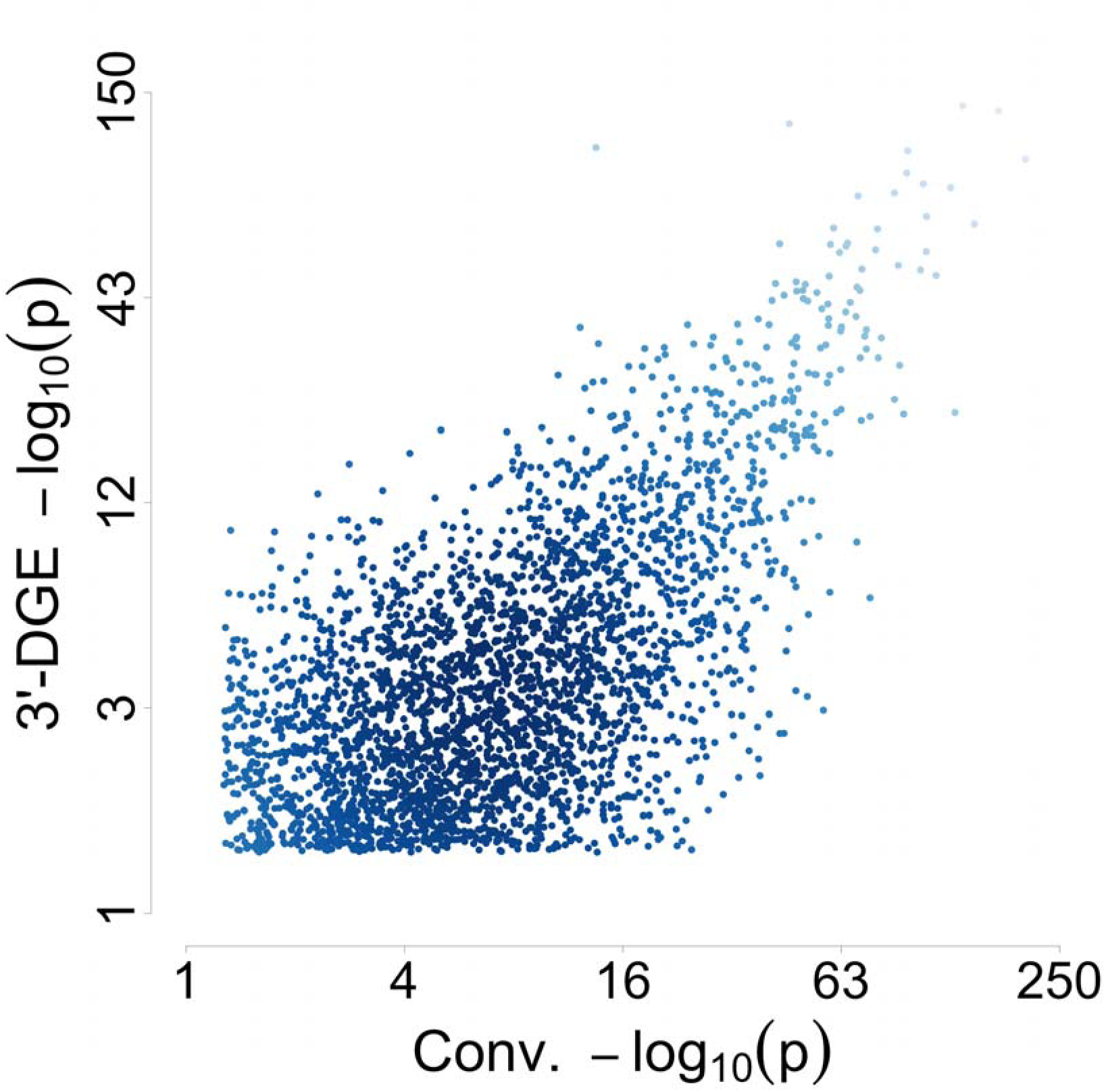
Comparison of Statistical Significance for the 3,136 Shared Differentially Expressed Genes (DEGs) from Sorafenib-treated Samples in 3’-end Digital Gene Expression (3’-DGE) and Conventional (Conv) Methods with FDR<0.1. The negative base-10 logarithm of the p-value for differential expression is plotted for each technique, with depth of color indicating density of points.

**Figure 5C.**
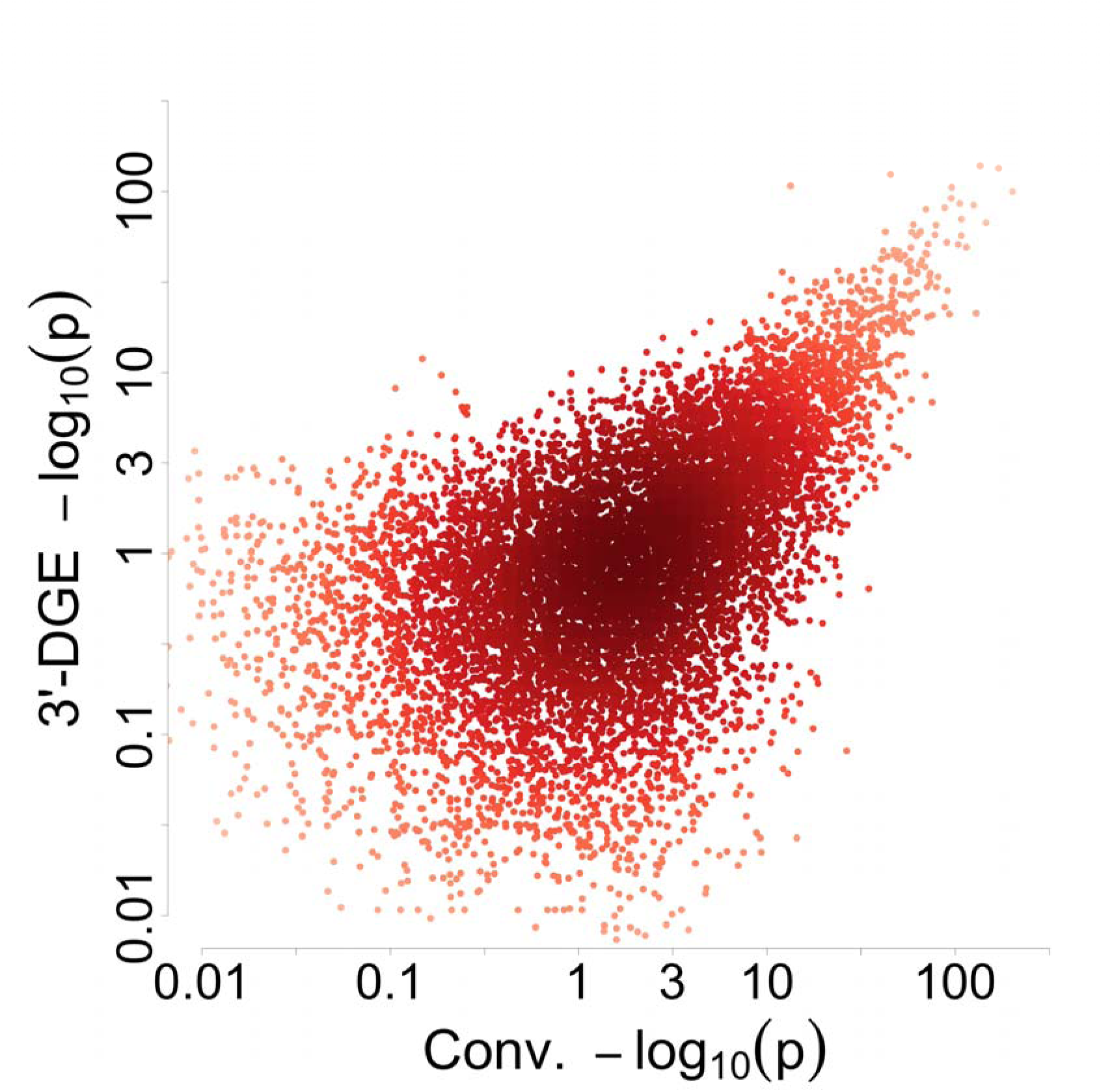
Comparison of Statistical Significance for All Genes Identified from Sorafenib-treated Samples by the 3’-end Digital Gene Expression (3’-DGE) and Conventional (Conv) Methods. The negative base-10 logarithm of the p-value for differential expression is plotted for each technique, with depth of color indicating density of points.

**Figure 5D.**
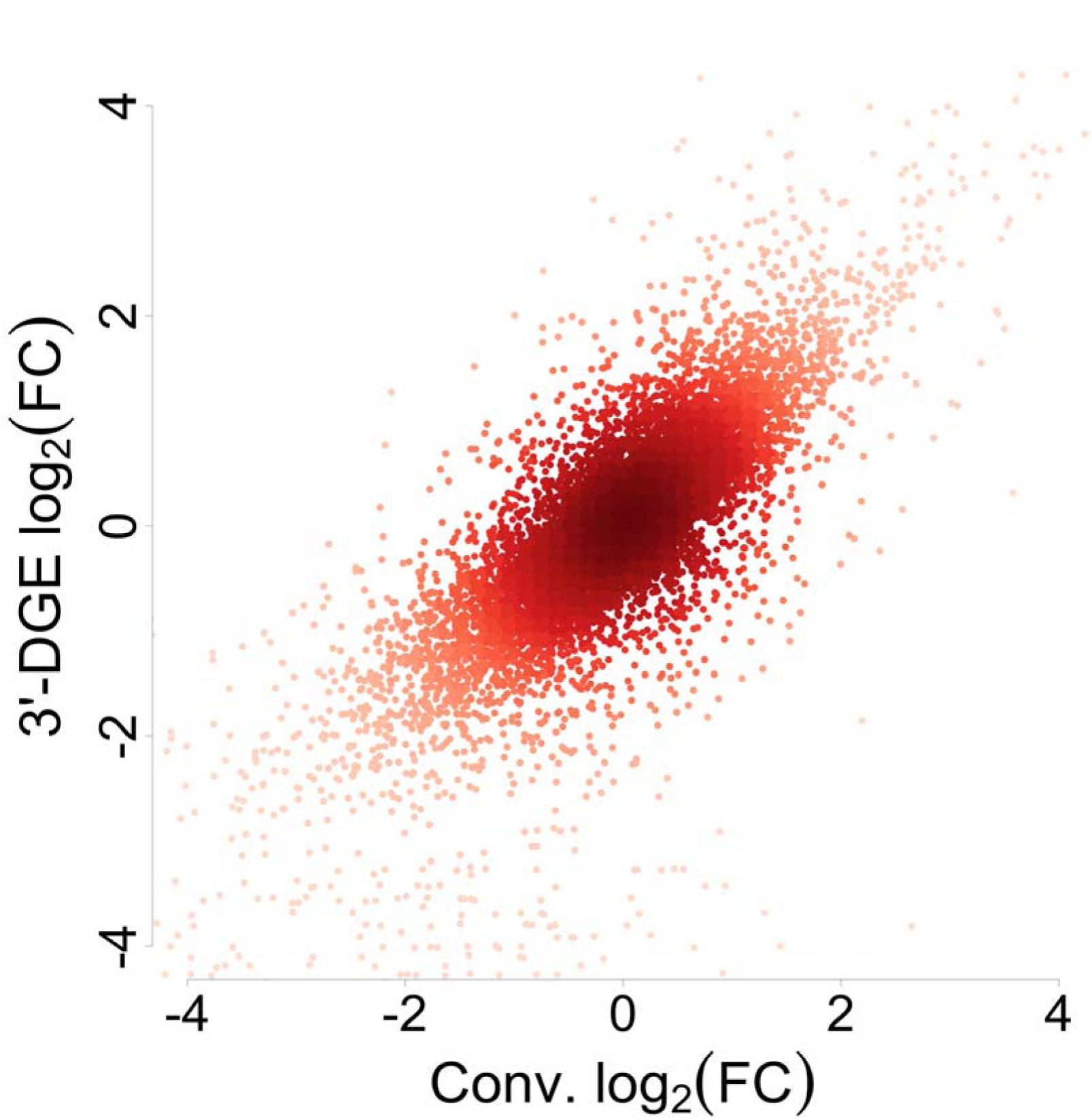
Comparison of Fold Change for All Genes Identified from Sorafenib-treated Samples by the 3’-end Digital Gene Expression (3’-DGE) and Conventional (Conv) Methods. The log base two fold-change is plotted for each technique, with depth of color indicating density of points.

**Figure 5E-F.**
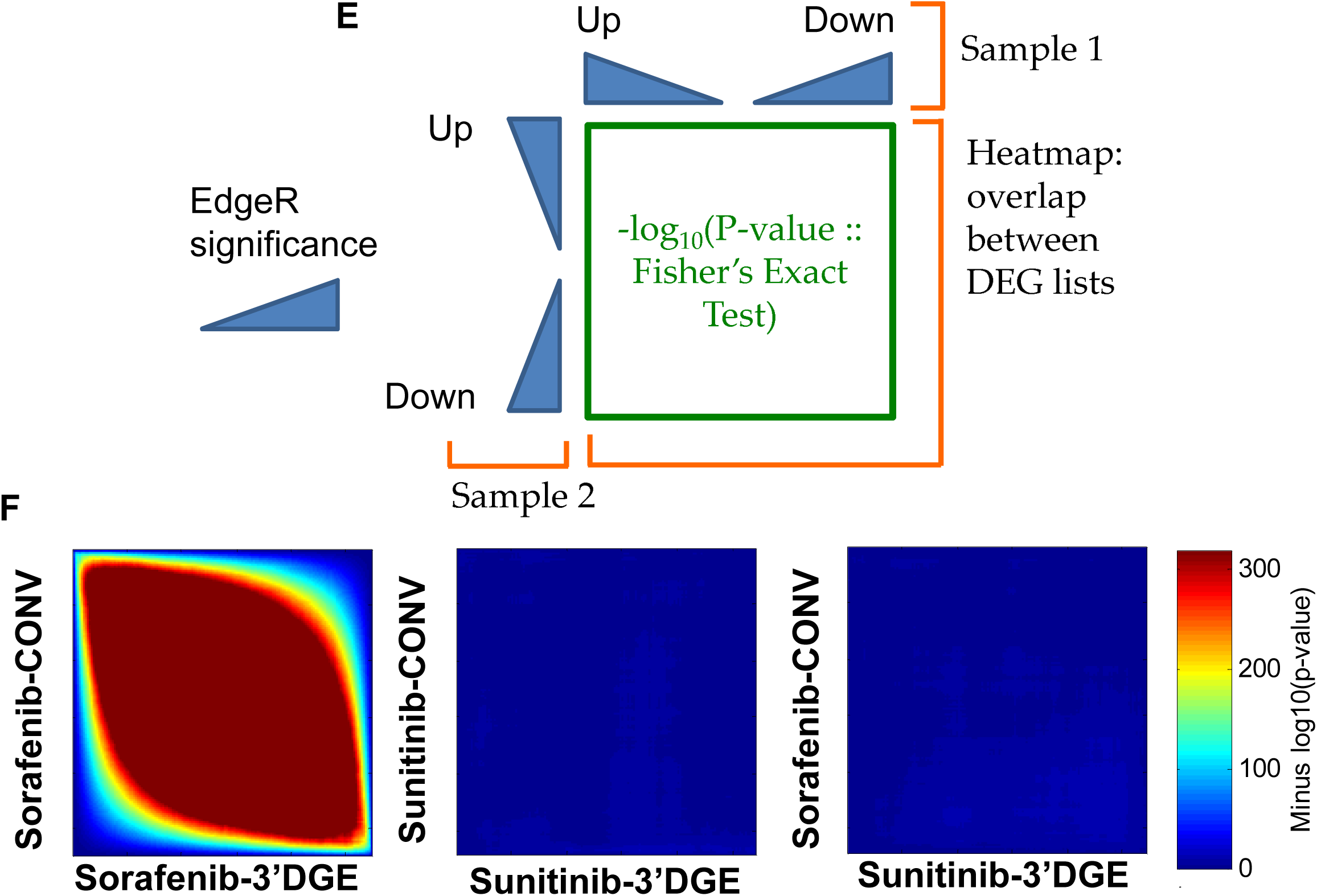
Rank-Rank Hypergeometric Tests for Consistency of Differential Expression Ranking and Gene Expression Signatures. **E.** All genes for which a p-value for differential expression was calculated were first sorted into up or down regulated genes (as compared to CTRL), and then ranked by statistical significance. The probability of overlap between two different such rank lists was calculated with Fisher’s Exact Test (aka hypergeometric test), and visualized with a heatmap, for all combinations of list cutoffs. **F.** Pairwise comparisons of SUN- and SOR-treated data for 3’-DGE and conventional. Only SOR-treated samples show significant overlap in their ranked lists, as expected, because only SOR induced large changes in gene expression. All other combinations have little evidence for statistical similarity in identified gene expression signatures.

### Comparison of 3’-DGE and Conventional with An Independent Dataset

Can we expect the agreement between 3’-DGE and conventional mRNAseq results to be similar for other datasets? We attempted to answer this question by taking a similar experimental design approach with a different set of RNA, this time from induced pluripotent stem cell (iPSC) lines created from control individuals or those that had a mutation for and clinical presentation of spinal muscular atrophy (SMA), where slightly different random primed library preparation and sequencing approaches were employed (see Methods). SMA is a childhood early onset motor neuron disease (Sareen et al. 2012; Fuller et al. 2015) where symptoms can be observed as early as 3 months or up to 2-3 years of age. The disease is typically caused by mutations in the SMN1 (Survival Motor Neuron 1) gene, however disease severity varies based on numbers of copies of the related SMN2 gene. As SMN genes control critical RNA biogenesis processes during early development (Fuller et al. 2015), we anticipate differential gene expression changes even at the iPSC stage without subsequent differentiation. For these analyses, two SMA subject clonal lines and three control lines, with varying numbers of growth replicates, were compared. Growth and disease replicates were combined for comparisons to generate the differentially expressed gene lists conventional mRNAseq and 3’DGEs. While this is not a traditional replicate approach for differential expression analysis, the mixed basis of the samples would only serve to increase sample-to-sample variability, and thus conclusions about consistency of differential expression can be considered as a lower bound that would only improve with other replication designs.

First, we analyzed sensitivity to detect gene expression as a function of read depth. We observed similar results as before, with conventional having higher sensitivity, detecting ~15% more genes (Fig. 6A), albeit now with both techniques having a higher level of overall detection as compared to the previous dataset. This could be due to different RNA preparation techniques used (TRIzol vs. Qiagen RNAeasy). We observed significant correlation between counts on a gene-by-gene level that again was greatly improved by normalizing conventional data by transcript lengths (Fig. 6B-C and Fig. S8). Because of the statistical power properties associated with UMI counting discussed above, it is not appropriate to look at strict FDR cutoffs to analyze the number of differentially expressed genes. Alternatively, we analyzed statistical significance (Fig. 6D) and log_2_ fold changes (Fig. 6E) for differential expression and found they were correlated well, and also similarly as before with sorafenib-treated samples. This leads to the resulting ranked gene signatures of differential expression again being highly concordant between the two techniques (Fig. 6F). Thus, we conclude that agreement reported here between 3’-DGE and conventional approaches is likely to be seen across many sample types, as well as across variations of conventional random primed methods.

**Figure 6A.**
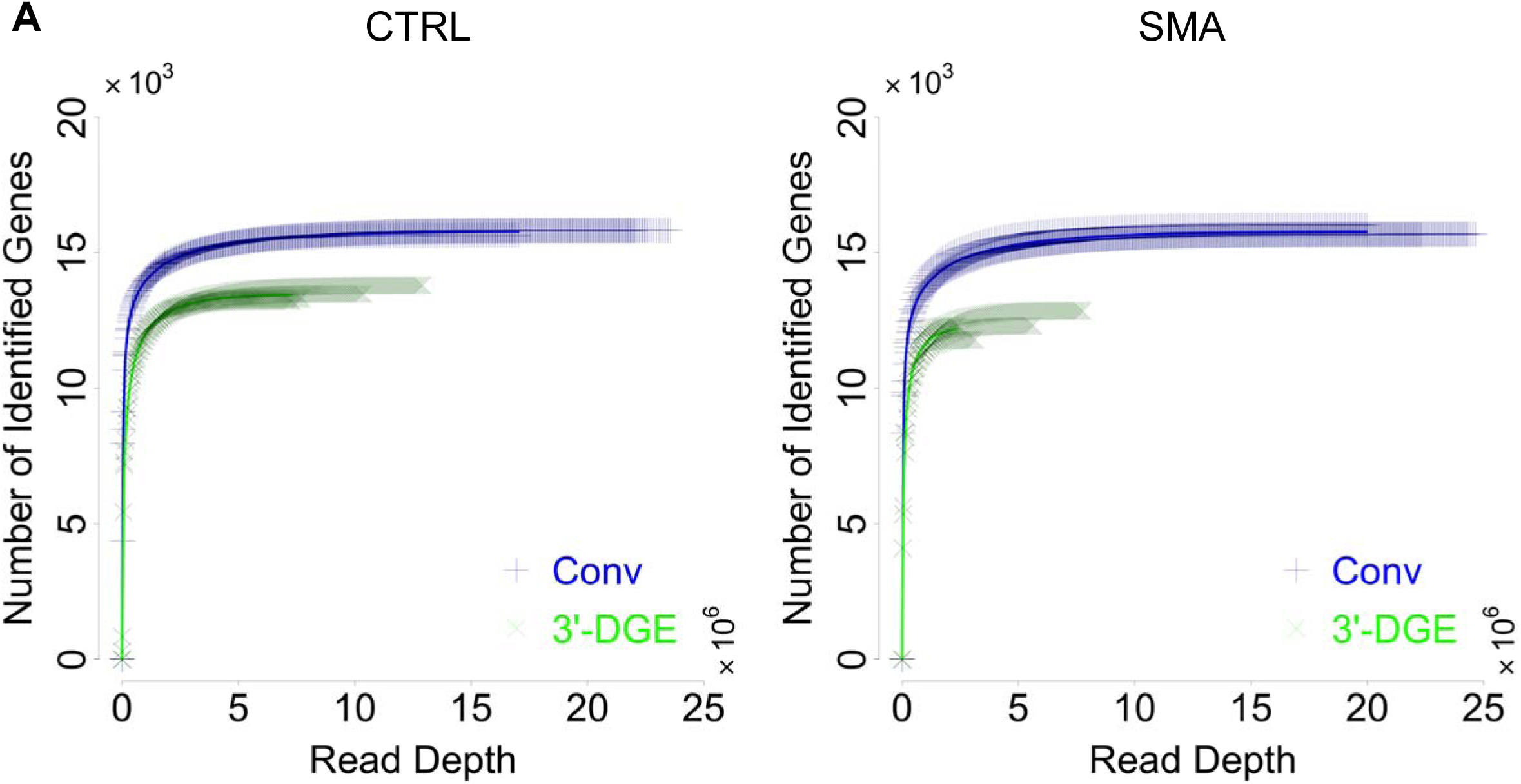
Sensitivity of Conventional (Conv) and 3’-end Digital Gene Expression (3’-DGE) mRNA Sequencing Methods in an Independent Dataset. Gene-wise reads are removed from every sample in a probability proportional to the abundance of the gene in a sample, to generate a set of the number of identified genes over a range of simulated read depths. The curves for individual replicate samples are shown with the thinner points, showing in general low variability. The average is shown with the solid line.

**Figure 6B-C.**
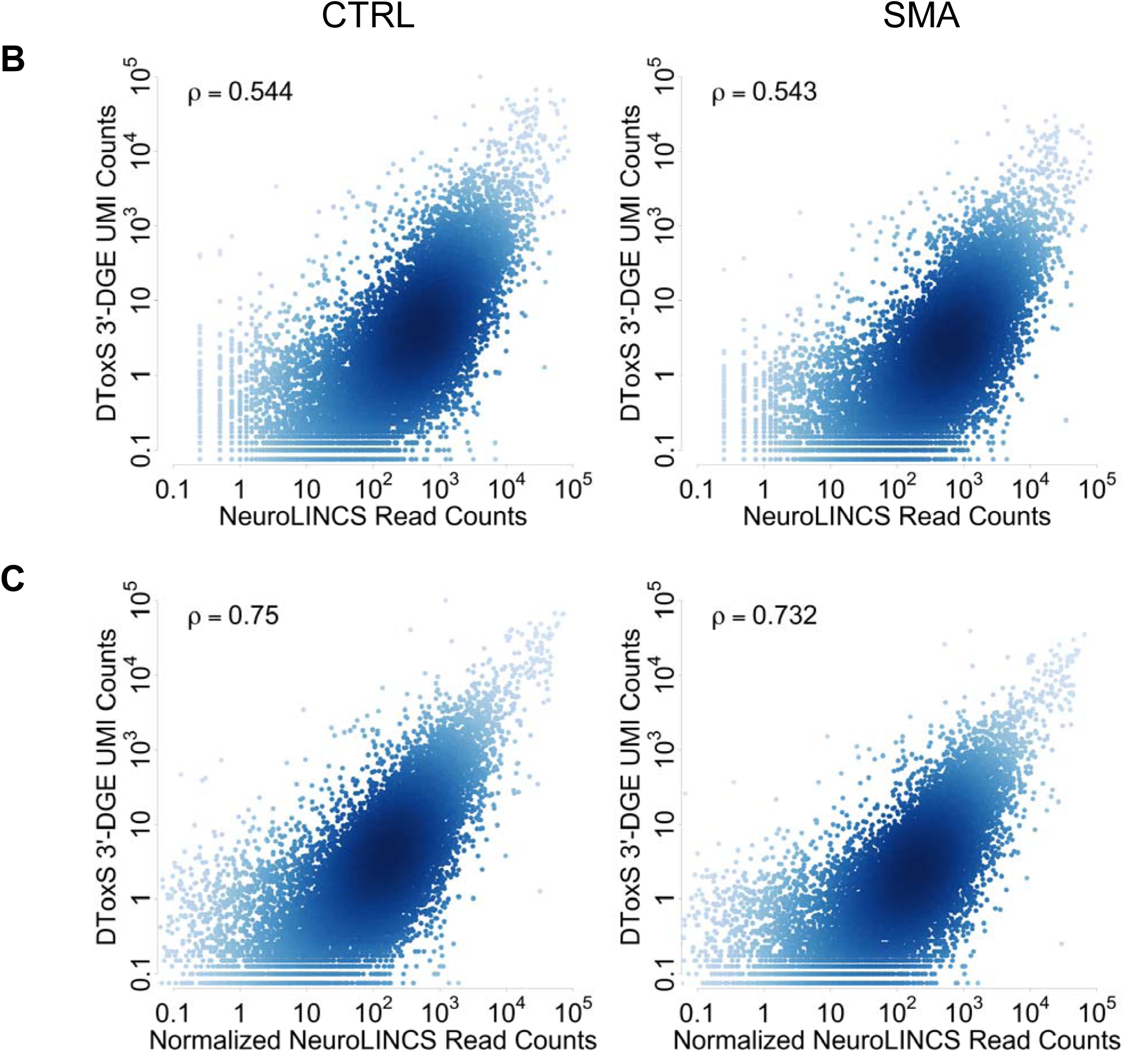
Quantitative Gene-wise Comparison between 3’-end Digital Gene Expression (3’-DGE) and for Independent Conventional Techniques. Density of points in scatter plots is indicated by depth of color. Inset text box shows Pearson correlation. In all plots, data are scaled so units are comparable. B. Scatterplots of UMI counts for DToxS’ 3’-DGE versus read counts for NeuroLINCS’ conventional, without normalization by average transcript length. CTRL or SMA refer to the genetic status of the iPS cells (see Methods). C. Scatterplots of UMI counts for DToxS’ 3’-DGE versus transcript length-normalized read counts for NeuroLINCS’ conventional.

**Figure 6D-E.**
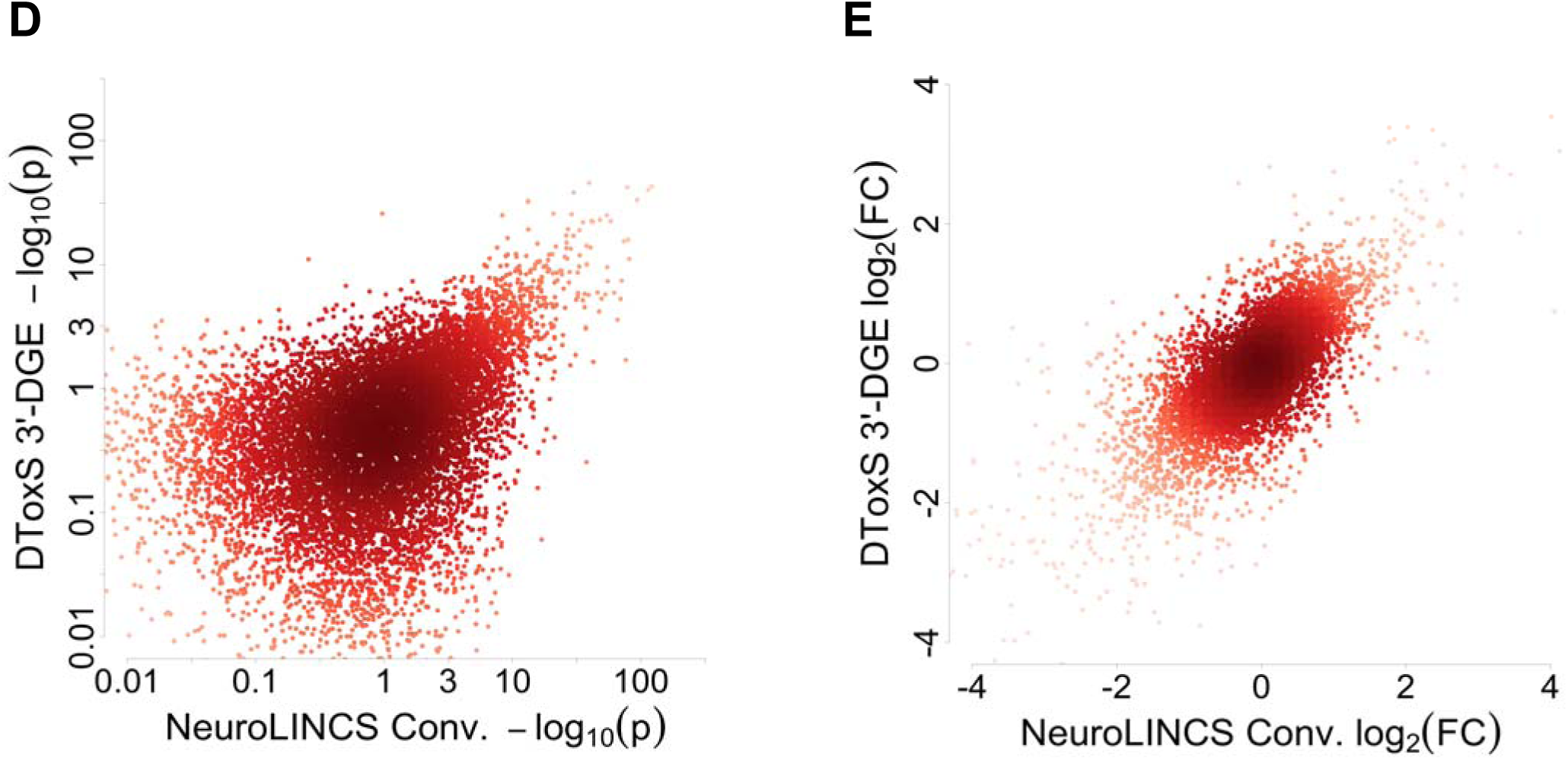
Comparison of Statistical Significance (D) or Fold-Change (E) for All Genes Identified from SMA Samples by DToxS’ 3’-end Digital Gene Expression (3’-DGE) and NeuroLINCS’ Conventional (Conv) Methods. **(D)** The negative base-10 logarithm of the p-value for differential expression is plotted for each technique, with depth of color indicating density of points. **(E)** The log base two fold-change is plotted for each technique, with depth of color indicating density of points.

**Figure 6F.**
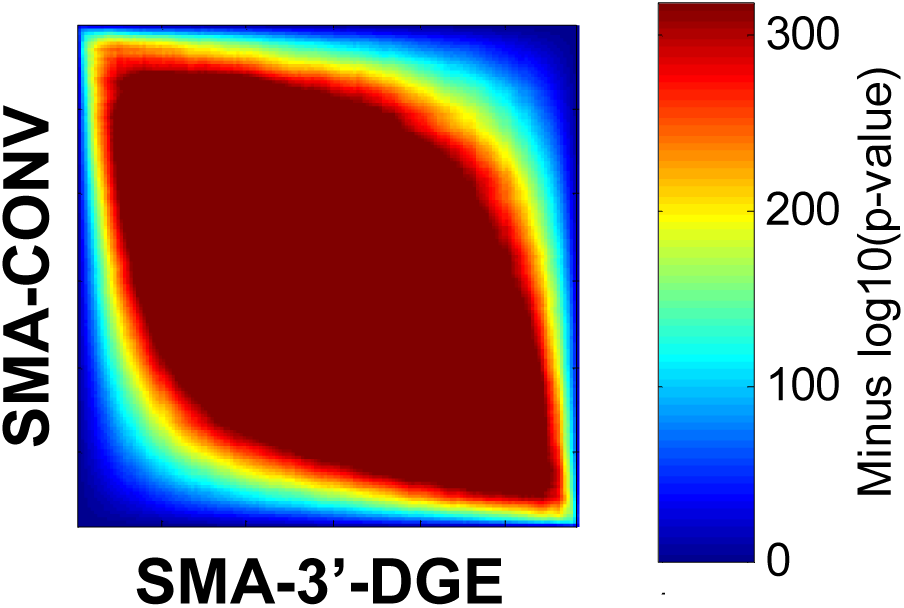
Rank-Rank Hypergeometric Tests for Consistency of Differential Expression Ranking and Gene Expression Signatures with an Independent Conventional Dataset. All genes for which a p-value for differential expression was calculated were first sorted into up or down regulated genes (as compared to CTRL), and then ranked by statistical significance. The probability of overlap between two different such rank lists was calculated with Fisher’s Exact Test (aka hypergeometric test), and visualized with a heatmap, for all combinations of list cutoffs. Shown here are lists from SMA vs. control for 3’-DGE and conventional.

## Methods

### Cell Culture and RNA Isolation

We used a commercially available cell line to compare the two sequencing methods. Detailed SOPs for culture and RNA isolation are available on our website (www.dtoxs.org) as DToxS SOP CE – 1.0: PromoCell Cardiomyocyte Subculture, DToxS SOP CE – 2.0: PromoCell Cardiomyocyte Plating for Drug Test, DToxS SOP CE – 4.0: Drug Treatment and Cell Lysis, and DToxS SOP A – 1.0: Total RNA Isolation.

Briefly, primary adult human cardiomyocytes (PromoCell, Heidelberg, Germany; Cat: C-12810, Lot: 3042901.2) were subcultured according to manufacturer’s instructions using antibiotic-free myocyte growth medium (PromoCell, Cat: C-22170) supplemented with a mix of 5% fetal calf serum, 0.5 ng/ml EGF, 2 ng/ml FGF, and 5 μg/ml recombinant human insulin (PromoCell, Cat: C-39275). We differentiated fully confluent cells for four weeks under serum starvation and treated cells with DMSO vehicle (Control, CTRL), 1 μM sorafenib (SOR), or 0.1 μM sunitinib (SUN) for 48 hours. Total RNA was extracted using TRIzol (Life Technologies, Cat: 15596018). RNA concentration was measured by Qubit 3.0 fluorometric quantitation using the high sensitivity RNA kit (Life Technologies, Cat: Q32852), and RNA quality was assessed by Fragment Analyzer using high sensitivity RNA analysis kit (Advanced Analytical Technologies, Cat: DNF-472). Only the RNA samples that have passed the quality control step with RNA integrity number (RIN) score of 7.0 or above were used. 200 ng of RNA for all samples were then transferred onto a skirted 96-well PCR plate (Eppendorf, Cat: 951020401) at a normalized concentration of 10 ng/μL.

### 3’-DGE Library Preparation

The RNA-seq libraries were prepared according to the Single Cell RNA Barcoding and Sequencing method originally developed for single cell RNA-seq (SCRB-seq; (Soumillon et al. 2014)) and adapted to extracted total RNA. Briefly, Poly(A)+ mRNA from extracted total RNA were converted to cDNA decorated with universal adapters, sample-specific barcodes and unique molecular identifiers (UMIs) using a template-switching reverse transcriptase. Decorated cDNA from multiple samples were then pooled, amplified and prepared for multiplexed sequencing using a modified transposon-based fragmentation approach that enriched for 3’ ends and preserved strand information. A detailed SOP is available at www.dtoxs.org (DToxS SOP A – 6.0: High-throughput mRNA Seq Library Construction for 3’ Digital Gene Expression (DGE))

### Conventional Random Primed Library Preparation

Conventional sequencing libraries were prepared using 200 ng of total RNA and the TruSeq RNA Library Prep Kit (Illumina, Cat: RS-122-2001) per manufacturer’s instructions, with mRNA was enriched via poly-A-selection using oligoDT beads.RNA was then thermally fragmented and converted to cDNA, adenylated for adaptor ligation and PCR amplified. Prior to sequencing, quality and concentration of the cDNA library was confirmed using Agilent Bioanalyzer and Qubit fluorometric quantitation.

### Sequencing using Illumina HiSeq Platform

Both the random primed and 3’-DGE libraries were sequenced using the Illumina HiSeq 2500 platform. cDNA libraries were loaded onto Illumina flowcells using the Illumina c-Bot, and conventional libraries were sequenced with 100 nucleotide paired-end reads per manufacturer’s instruction, whereas 3’-DGE used a custom paired end protocol with 26 bp on the first read, and 46 on the second. Detailed SOPs for the 3’-DGE sequencing are available on www.dtoxs.org as described above.

### iPSC Cell Culture, RNA Isolation and Conventional Sequencing with iPSC Data

Human iPSCs were generated using episomal reprogramming and validated using quality control methods, including Pluritest, G-band karyotype analysis and Short Tandem Repeat identity test assays as described previously (Fuller et al. 2015). iPSCs were maintained in 6-well tissue culture plates coated with Corning Growth Factor Reduced Matrigel Matrix (Cat # 354230). iPSCs were passaged every 7 days or when the cultures reached 80-90% confluency using the StemPro EZPassaging Tool (Life Technologies, Cat # 23181010). To prepare cells for cell pellets, each iPSC line was seeded into two 6-well plates at a 1:6 dilution and allowed to reach ~80% confluency (approximately 1.5 to 2e6 cells per well). On the day of collection, any visible spontaneous differentiation was removed using a pipette tip. The spent media was then aspirated and the cells were rinsed with chilled PBS. 1ml of chilled PBS was added to each well and the cells were lifted gently from the plate using a Corning cell scraper (Cat # CC3010). Two wells from each plate were pooled together into a chilled 15ml conical and labeled as replicate #1. This was repeated until three replicates were collected per sample. Each replicate was then evenly distributed into 5 chilled 0.5ml Eppendorf tubes. The Eppendorf tubes were labeled with the cell line name, passage number, collection date and replicate number. Cells were then pelleted by centrifuging each sample for 5 minutes at 1000 rpm. The PBS was aspirated and the cell pellets were then flash frozen by briefly submerging each tube in liquid nitrogen. The samples were sent for RNA preps and sequencing. Full details relating to Control and SMA cell lines are also available at http://lincsportal.ccs.miami.edu/dcic-portal/.

Total RNA were isolated using the Qiagen RNeasy Kit and QIAshredder. RNA QC were analyzed on the Agilent 2100 Bioanalyzer which indicated that all RIN values were 10. RNA-Seq libraries were made with 1 ug of RNA using the Illumina non-stranded TruSeq mRNA v2 protocol. Libraries were quantified using the KAPA library quant kit and sequenced on the HiSeq 2500 using 100 cycles across three lanes to obtain paired-end reads 100 base pairs in length.

### Defining and Alignment to Reference Genome

For the conventional method, we obtained the genome sequence file (hg19) from the UCSC website (http://hgdownload.cse.ucsc.edu/downloads.html#human). We created the gene annotation file by downloading a list of RefSeq genes from *Table Browser* (http://genome.ucsc.edu/cgi-bin/hgTables) and then converting it to the GTF format using *genePredToGtf* (http://hgdownload.cse.ucsc.edu/admin/exe). For 3’-DGE, we used a shortened version of hg19 limited to smaller regions around the 3’ ends of genes (available upon request). There are a common set of 22,081 genes which we used as a basis of comparison between the two techniques throughout the manuscript (Table S1). http://genome.ucsc.edu/cgi-bin/hgTables) and then converting it to the GTF format using *genePredToGtf* (http://hgdownload.cse.ucsc.edu/admin/exe). For 3’-DGE, we used a shortened version of hg19 limited to smaller regions around the 3’ ends of genes (available upon request). There are a common set of 22,081 genes which we used as a basis of comparison between the two techniques throughout the manuscript (Table S1). We used STAR (Dobin et al. 2013) with the default parameter settings to align the conventional mRNAseq data (PromoCell) to the reference described above, and then counted the number of the sequence fragments uniquely aligned to each gene by the *featureCounts* program from a sequence alignment suite *Subread* using the UCSC reference gene annotation. For 3’-DGE data, a custom python script is used (available upon request). First, reads are aligned using BWA, and counts of specifically or non-specifically aligned reads (i.e., aligning to one or more than one gene with high confidence) are calculated. Next, the number of distinct unique molecular identifier (UMI) sequences embedded in those aligned reads are counted, giving the UMI counts.

### Computational Downsampling of Sequencing Depth

To compare datasets on an equivalent sequencing depth basis, we computationally removed read counts with an iterative algorithm (Fig. S4-S5). First, all genes with very low expression are removed (<4 read counts). Then, a particular gene is randomly selected with probability proportional to its count representation amongst all genes, and a read is removed from this gene. We removed UMI counts from the selected gene probabilistically, according to the ratio of UMI counts to read counts (always less than one, and a gene-specific estimate of PCR bias). The process is repeated using probabilities estimated at the first iteration until all read counts are removed. We performed the process 16 times for each sample to ensure the stochastic nature of removal did not affect our results (Fig. S6).

### Data Normalization

For conventional data, we divided read counts by the average transcript length. These lengths were obtained from http://genome.ucsc.edu/cgi-bin/hgTables, by summing the length of all exons in a transcript into a transcript length, and then averaging this transcript length across all transcripts of each gene.

### Differential Expression Analysis

We performed differential expression analysis for the PromoCell data with edgeR (Robinson et al. 2009) starting with tables of counts for any technique. Differential expression analysis consists of the following steps within edgeR: normalization by trimmed mean of M-values (Robinson and Oshlack 2010), empirical Bayes estimation of sample dispersion (Chen et al. 2014), and exact test for negative-binomial sample comparison (Robinson and Smyth 2008). A detailed standard operating procedure (SOP) document is available at www.dtoxs.org, and all code is available upon request.

### Rank-rank hypergeometric test

The significance of the overlap between two lists of differentially expressed genes (DEGs) was calculated using the rank-rank hypergeometric test (Plaisier et al. 2010). First, for all genes, we calculated the negative log10 of the p-value for differential expression (−log10*p*), and multiplied by negative one if a gene was downregulated. Genes were ranked by signed −log_10_*p*, placing the most significantly up-regulated genes at the top and the most significantly down-regulated genes at the bottom of the list. The number of overlapping genes of the top *x* genes of one list and the top *y* genes of the other was counted at every 10^th^ *x-y* combination, and Fisher’s exact test was used to calculate significance of the overlap. If the overlap was greater than expected the right-tailed Fisher’s Exact test was used, otherwise the left-tailed Fisher’s exact test was used. The resulting heatmap of p-values from Fisher’s Exact text was visualized with the MATLAB function `imagesc`.

## Supplementary Figures

**Figure S1.**
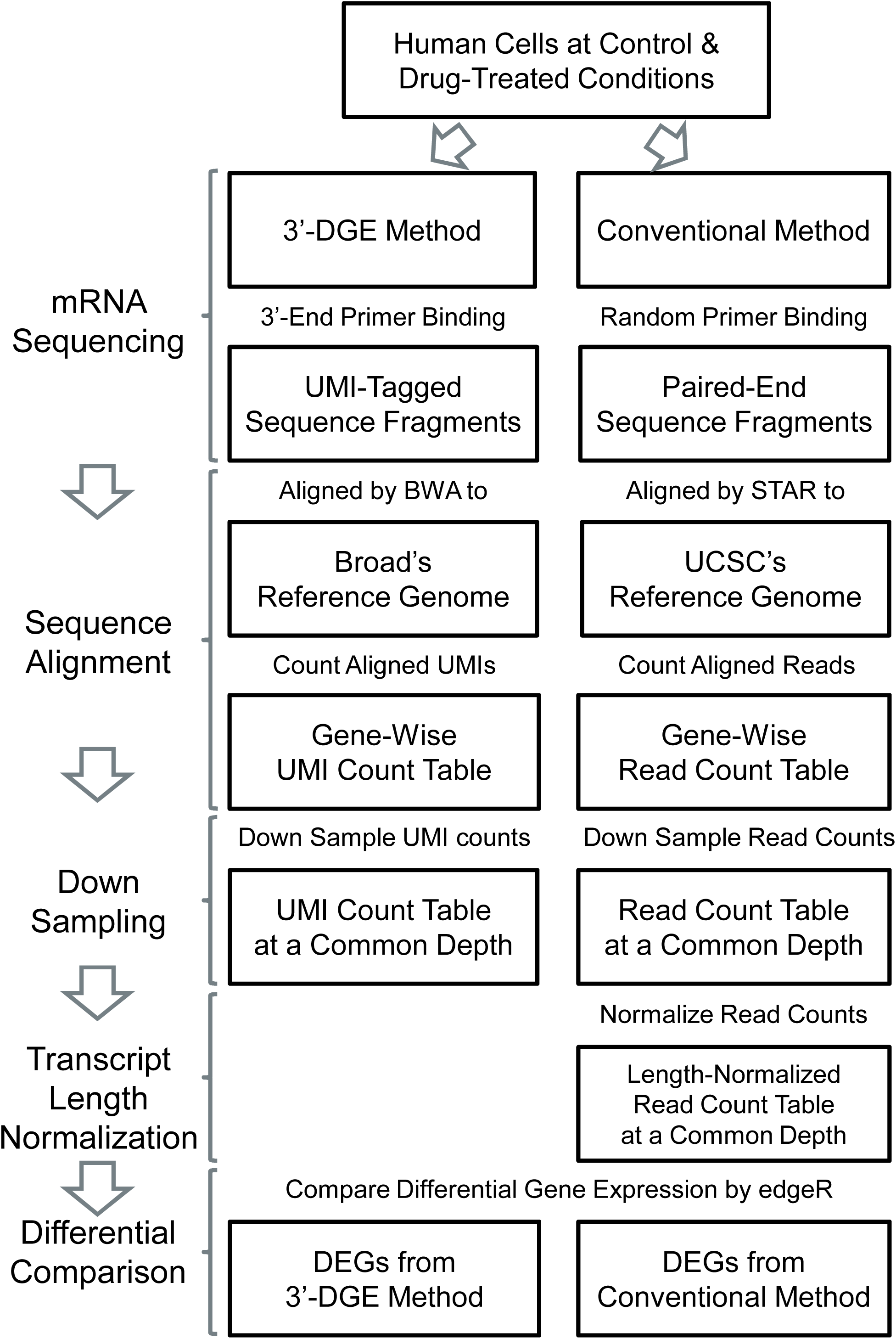
The Workflow of Comparing Two mRNA Sequencing Methods: the 3’-end Digital Gene Expression (3’-DGE) Method and the Conventional Random Primer-binding Method.

**Figure S2.**
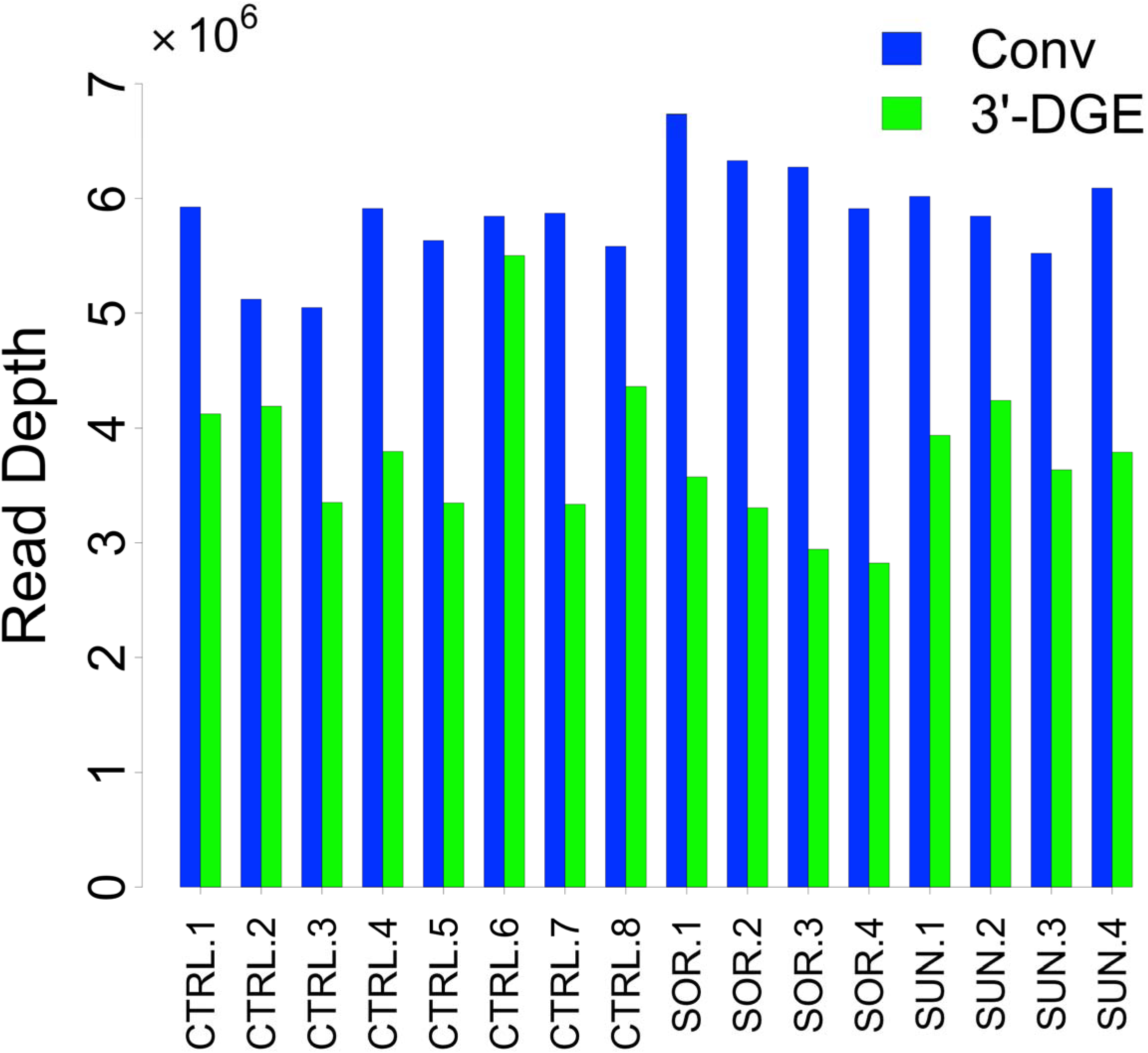
Read Depth Across Multiple Samples for Conventional (Conv) and 3’-end Digital Gene Expression (3’-DGE) Methods. The total number of uniquely aligned reads is plotted for each sample across the three treatment conditions: control (DMSO-CTRL), Sorafenib (SOR), and Sunitinib (SUN)). All samples show consistent read depth.

**Figure S3.**
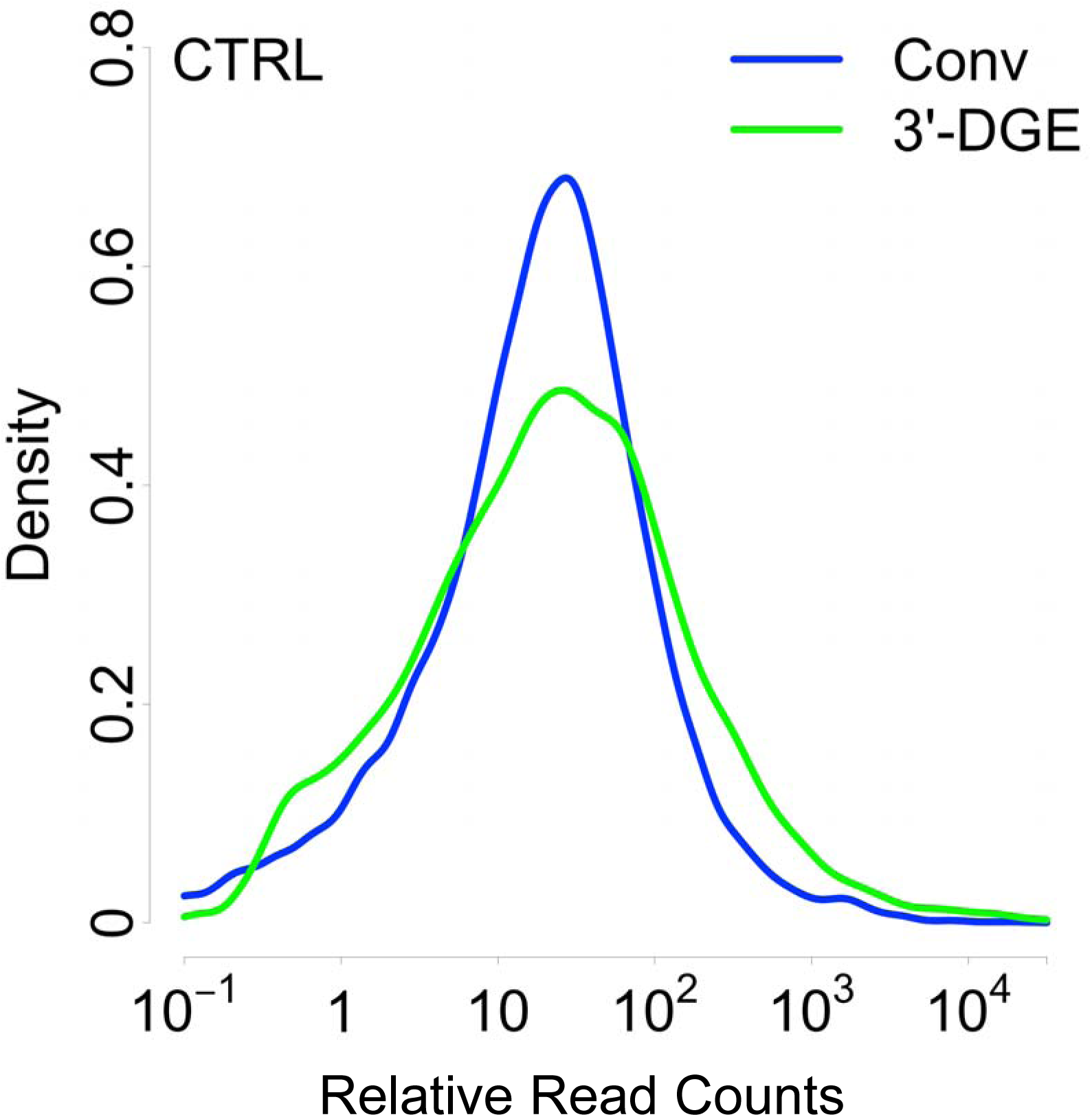
Relative Read Count Distributions for Conventional and 3’-DGE Methods. The mean read counts for each gene across the eight control samples, downsampled to a common read depth (2.8 million per sample) was calculated, and for conventional, this value was divided by transcript length. The probability density was estimated by the *density* function of R package *stats*.

**Figure S4.**
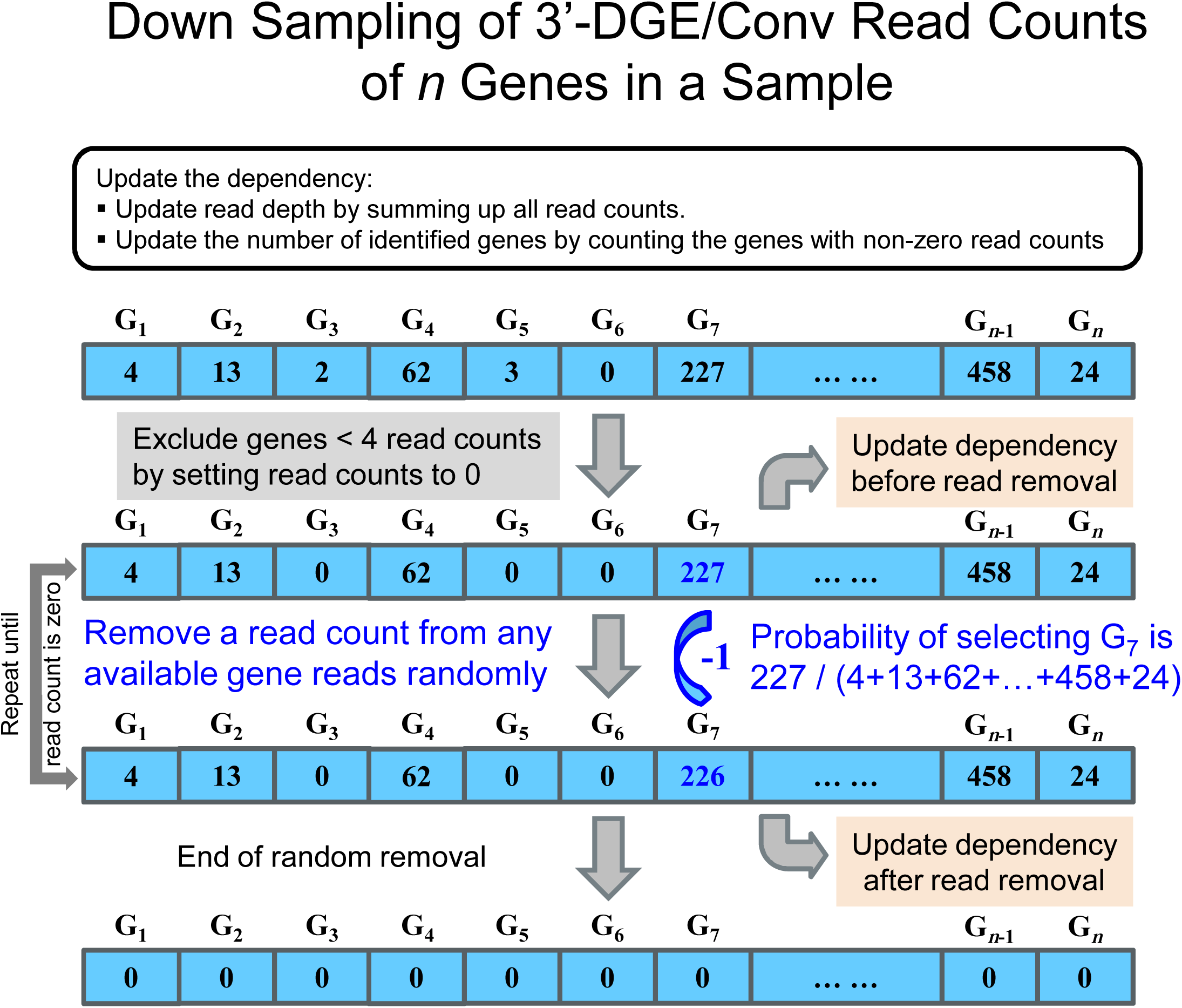
Down-sampling Read Counts in 3’-end Digital Gene Expression (3’-DGE) and Conventional (Conv) Sequencing Methods.

**Figure S5.**
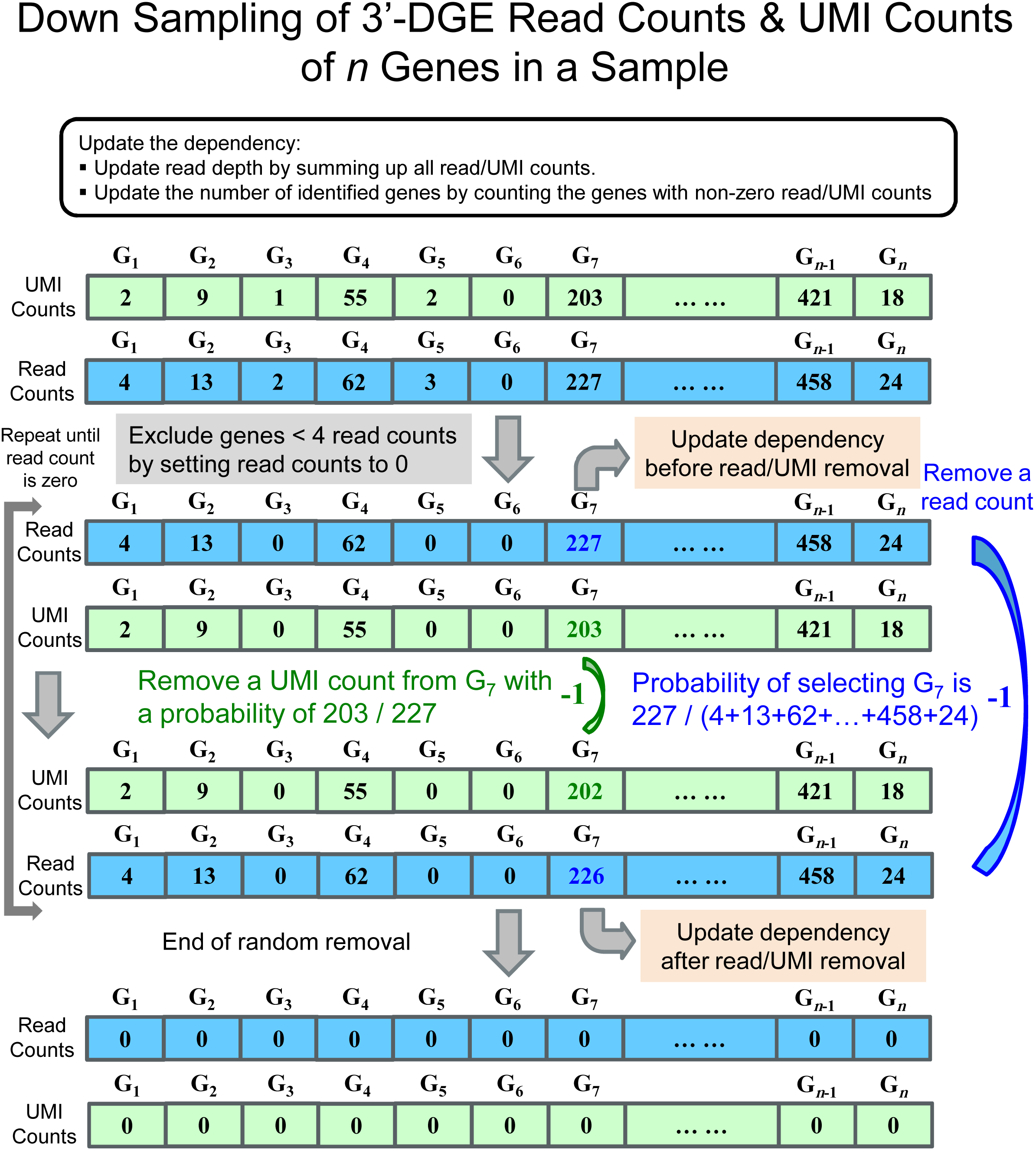
Down-sampling Read Counts and Unique Molecular Identifier Counts for the 3’-Digital Gene Expression (3’-DGE) Method.

**Figure S6.**
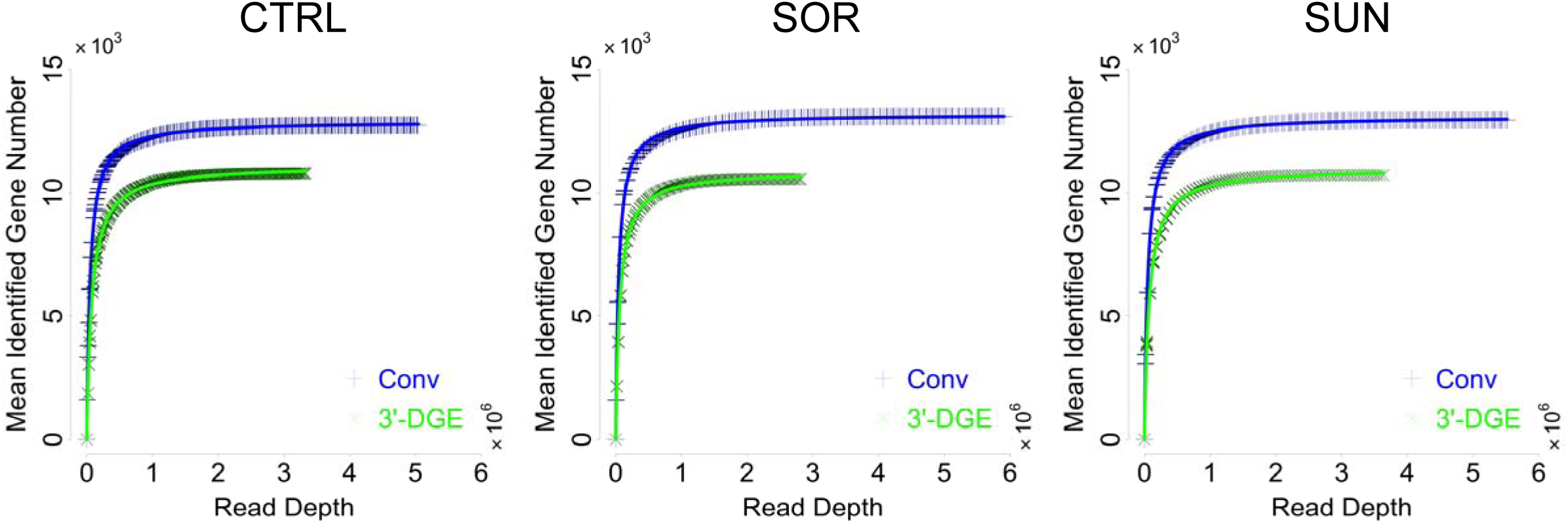
Variability in the Read Removal Process. Random read removal was performed 16 independent times, and the range of variability across those runs is not visible on this chart despite 16 different runs being plotted, indicating a highly reproducible simulation algorithm for read removal on the level of identified genes.

**Figure S7.**
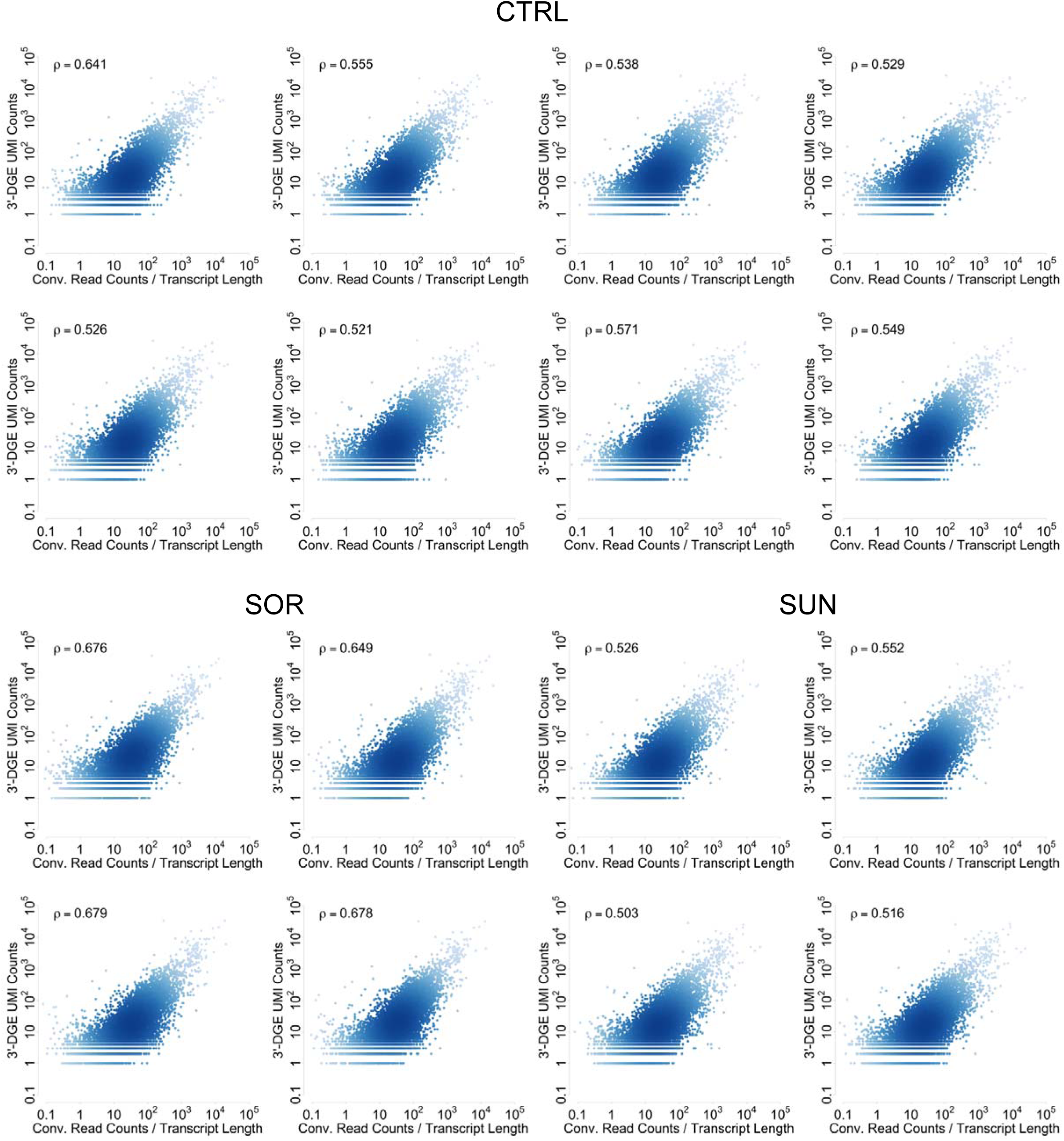
Sample-to-Sample Comparison between Two Techniques. Datasets are down-sampled to a common read depth of 2.8 million reads, and then gene-by-gene comparisons are made via scatter plots. To generate a reduced UMI count dataset, upon removal of a read count, UMI counts were removed with probability proportional to the ratio between UMI counts and read counts for that gene (accounting for PCR bias). Density of points in scatter plots is indicated by depth of color. Inset text box shows Pearson correlation. In all plots, data are scaled so units are comparable. There are eight CTRL samples, four SOR samples, and four SUN samples. All are biological replicates.

**Figure S8.**
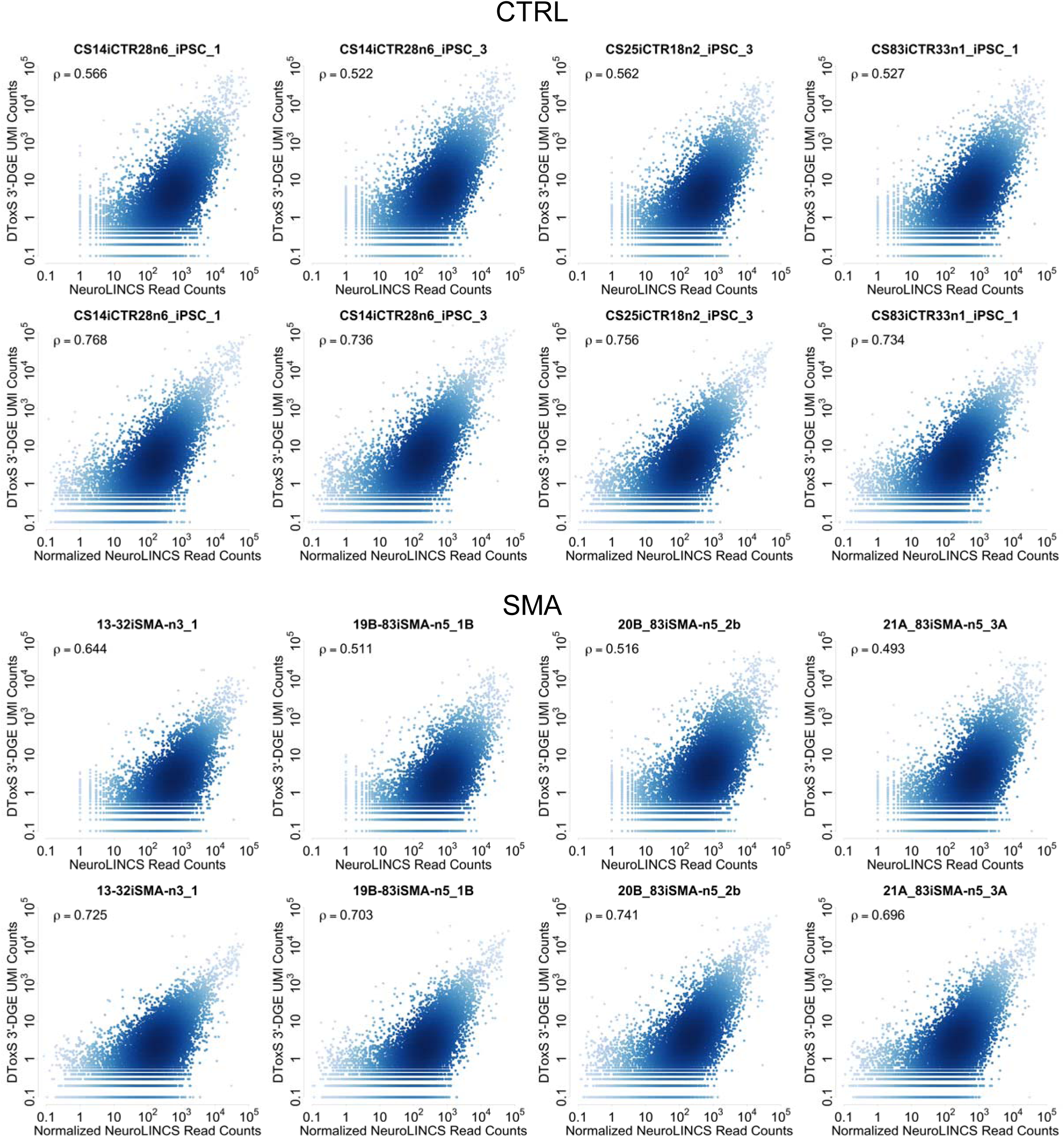
Sample-to-Sample Comparison between 3’-end Digital Gene Expression (3’-DGE) and Independent Conventional Techniques. Density of points in scatter plots is indicated by depth of color. Inset text box shows Pearson correlation. In all plots, data are scaled so units are comparable. There are four CTRL samples and four SMA samples. All are biological replicates.

## References

Ardlie KG, Deluca DS, Segre A V., Sullivan TJ, Young TR, Gelfand ET, Trowbridge CA, Maller JB, Tukiainen T, Lek M, et al. 2015. The Genotype-Tissue Expression (GTEx) pilot analysis: Multitissue gene regulation in humans. Science (80-) 348: 648–660. http://www.sciencemag.org/content/348/6235/648.full.

Byron SA, Van Keuren-Jensen KR, Engelthaler DM, Carpten JD, Craig DW. 2016. Translating RNA sequencing into clinical diagnostics: opportunities and challenges. Nat Rev Genet 17: 257–271. http://www.nature.com/doifinder/10.1038/nrg.2016.10%5Cn http://dx.doi.org/10.1038/nrg.2016.10.

Chen Y, Lun ATL, Smyth GK. 2014. Differential Expression Analysis of Complex RNA-seq Experiments Using edgeR. Stat Anal Next Gener Seq Data 1–25.

Conesa A, Madrigal P, Tarazona S, Gomez-Cabrero D, Cervera A, McPherson A, Szcześniak MW, Gaffney DJ, Elo LL, Zhang X, et al. 2016. A survey of best practices for RNA-seq data analysis. Genome Biol 17: 13. http://genomebiology.biomedcentral.com/articles/10.1186/s13059-016-0881-8.

DeLaughter DM, Bick AG, Wakimoto H, McKean D, Gorham JM, Kathiriya IS, Hinson JT, Homsy J, Gray J, Pu W, et al. 2016. Single-Cell Resolution of Temporal Gene Expression during Heart Development. Dev Cell 0: 163–172. http://linkinghub.elsevier.com/retrieve/pii/S1534580716306827.

Dobin A, Davis CA, Schlesinger F, Drenkow J, Zaleski C, Jha S, Batut P, Chaisson M, Gingeras TR. 2013. STAR: ultrafast universal RNA -seq aligner. Bioinformatics 1–7.

Fuller HR, Mandefro B, Shirran SL, Gross AR, Kaus AS, Botting CH, Morris GE, Sareen D. 2015. Spinal Muscular Atrophy Patient iPSC-Derived Motor Neurons Have Reduced Expression of Proteins Important in Neuronal Development. Front Cell Neurosci 9: 506. http://www.pubmedcentral.nih.gov/articlerender.fcgi?artid=4707261&tool=pmcentrez&rendertype=abstract.

Genomics H, Rivas MA, Pirinen M, Conrad DF, Lek M, Tsang EK, Karczewski KJ, Maller JB, Kukurba KR, Deluca DS, et al. 2015. COHORT. 348. http://science.sciencemag.org/content/sci/348/6235/666.full.pdf.

Islam S, Zeisel A, Joost S, La Manno G, Zajac P, Kasper M, Lönnerberg P, Linnarsson S. 2014. Quantitative single-cell RNA-seq with unique molecular identifiers. Nat Methods 11: 163–166. http://www.ncbi.nlm.nih.gov/pubmed/24363023.

Kivioja T, Vähärautio A, Karlsson K, Bonke M, Enge M, Linnarsson S, Taipale J. 2011. Counting absolute numbers of molecules using unique molecular identifiers. Nat Methods 9: 72–74. http://www.nature.com/nmeth/journal/v9/n1/pdf/nmeth.1778.pdf.

La Manno G, Gyllborg D, Codeluppi S, Nishimura K, Salto C, Zeisel A, Borm LE, Stott SRW, Toledo EM, Villaescusa JC, et al. 2016. Molecular Diversity of Midbrain Development in Mouse, Human and Stem Cells. Cell (in press). http://ac.els-cdn.com/S0092867416313095/1-s2.0-S0092867416313095-main.pdf?_tid=36fb8d98-b0f0-11e6-a01a-00000aacb35f&acdnat=1479845824_04308cb93aa0d4e8c1bceb46aa85c6b6.

Li S, Labaj PP, Zumbo P, Sykacek P, Shi W, Shi L, Phan J, Wu P-Y, Wang M, Wang C, et al. 2014a. Detecting and correcting systematic variation in large-scale RNA sequencing data. Nat Biotechnol 32: 888–895. http://eutils.ncbi.nlm.nih.gov/entrez/eutils/elink.fcgi?dbfrom=pubmed&id=25150837&retmode=ref&cmd=prlinks%5Cnpapers3://publication/doi/10.1038/nbt.3000.

Li S, Tighe SW, Nicolet CM, Grove D, Levy S, Farmerie W, Viale A, Wright C, Schweitzer PA, Gao Y, et al. 2014b. Multi-platform assessment of transcriptome profiling using RNA-seq in the ABRF next-generation sequencing study. Nat Biotechnol 32: 915–925. http://dx.doi.org/10.1038/nbt.2972%5Cn http://www.nature.com/doifinder/10.1038/nbt.2972

Melé M, Ferreira PG, Reverter F, Deluca DS, Monlong J, Sammeth M, Young TR, Goldmann JM, Pervouchine DD, Sullivan TJ, et al. 2015. Tissues and Individuals. Science (80-) 348: 660–665.

Mortazavi A, Williams BA, McCue K, Schaeffer L, Wold B. 2008. Mapping and quantifying mammalian transcriptomes by RNA-Seq. Nat Methods 5: 621–628. http://dx.doi.org/10.1038/nmeth.1226%5Cn http://www.nature.com/nmeth/journal/v5/n7/suppinfo/nmeth.1226_S1.html%5Cn http://www.nature.com/doifinder/10.1038/nmeth.1226%5Cn http://www.ncbi.nlm.nih.gov/pubmed/18516045.

Olsson A, Venkatasubramanian M, Chaudhri VK, Aronow BJ, Salomonis N, Singh H, Grimes HL. 2016. Single-cell analysis of mixed-lineage states leading to a binary cell fate choice. Nature 537: 1–22. http://www.nature.com/doifinder/10.1038/nature19348.

Plaisier SB, Taschereau R, Wong JA, Graeber TG. 2010. Rank-rank hypergeometric overlap: identification of statistically significant overlap between gene-expression signatures. Nucleic Acids Res 38: e169.

Risso D, Ngai J, Speed TP, Dudoit S. 2014. Normalization of RNA-seq data using factor analysis of control genes or samples. Nat Biotech 32: 896–902. http://www.pubmedcentral.nih.gov/articlerender.fcgi?artid=4404308&tool=pmcentrez&rendertype=abstract.

Robinson M, Oshlack A. 2010. A scaling normalization method for differential expression analysis of RNA-seq data. Genome Biol 11: R25. papers://08004521-e080-41da-b763-67c892855688/Paper/p391.

Robinson MD, McCarthy DJ, Smyth GK. 2009. edgeR: A Bioconductor package for differential expression analysis of digital gene expression data. Bioinformatics 26: 139–140. https://www.ncbi.nlm.nih.gov/pmc/articles/PMC2796818/pdf/btp616.pdf.

Robinson MD, Smyth GK. 2008. Small-sample estimation of negative binomial dispersion, with applications to SAGE data. Biostatistics 9: 321–332.

Sareen D, Ebert AD, Heins BM, McGivern J V., Ornelas L, Svendsen CN. 2012. Inhibition of apoptosis blocks human motor neuron cell death in a stem cell model of spinal muscular atrophy. PLoS One 7.

Smith GR, Birtwistle MR. 2016. A mechanistic beta-binomial probability model for mRNA sequencing data. PLoS One 11.

Soumillon M, Cacchiarelli D, Semrau S, van Oudenaarden A, Mikkelsen TS. 2014. Characterization of directed differentiation by high-throughput single-cell RNA-Seq. bioRxiv 3236. http://biorxiv.org/content/early/2014/03/05/003236.abstract.

Su Z, Łabaj PP, Li SS, Thierry-Mieg J, Thierry-Mieg D, Shi W, Wang C, Schroth GP, Setterquist R a, Thompson JF, et al. 2014. A comprehensive assessment of RNA-seq accuracy, reproducibility and information content by the Sequencing Quality Control Consortium. Nat Biotechnol 32: 903–14. http://www.pubmedcentral.nih.gov/articlerender.fcgi?artid=4321899&tool=pmcentrez&rendertype=abstract%5Cn http://www.nature.com/doifinder/10.1038/nbt.2957.

Trapnell C, Hendrickson DG, Sauvageau M, Goff L, Rinn JL, Pachter L. 2013. Differential analysis of gene regulation at transcript resolution with RNA-seq. Nat Biotechnol 31: 46–53. http://dx.doi.org/10.1038/nbt.2450.

Trapnell C, Pachter L, Salzberg SL. 2009. TopHat: Discovering splice junctions with RNA-Seq. Bioinformatics 25: 1105–1111. https://www.ncbi.nlm.nih.gov/pmc/articles/PMC2672628/pdf/btp120.pdf.

Trapnell C, Williams BA, Pertea G, Mortazavi A, Kwan G, van Baren MJ, Salzberg SL, Wold BJ, Pachter L. 2010. Transcript assembly and quantification by RNA-Seq reveals unannotated transcripts and isoform switching during cell differentiation. Nat Biotechnol 28: 511–515. http://www.nature.com/nbt/journal/v28/n5/full/nbt.1621.html%5Cn http://www.nature.com/nbt/journal/v28/n5/pdf/nbt.1621.pdf.

Wagner A, Regev A, Yosef N. 2016. Uncovering the vectors of cellular states with single cell genomics. Nat Publ Gr 34: 1–53. http://dx.doi.org/10.1038/nbt.3711.

Wang C, Gong B, Bushel PR, Thierry-Mieg J, Thierry-Mieg D, Xu J, Fang H, Hong H, Shen J, Su Z, et al. 2014. The concordance between RNA-seq and microarray data depends on chemical treatment and transcript abundance. Nat Biotechnol 32: 926–32. http://www.pubmedcentral.nih.gov/articlerender.fcgi?artid=4243706&tool=pmcentrez&rendertype=abstract.

